# Transcriptional and chromatin-based partitioning mechanisms uncouple protein scaling from cell size

**DOI:** 10.1101/2020.08.28.272690

**Authors:** Matthew P. Swaffer, Devon Chandler-Brown, Jacob Kim, Maurice Langhinrichs, Georgi Marinov, William Greenleaf, Anshul Kundaje, Kurt M. Schmoller, Jan M. Skotheim

## Abstract

Biosynthesis scales with cell size such that protein concentrations generally remain constant as cells grow. As an exception, synthesis of the cell-cycle inhibitor Whi5 ‘sub-scales’ with cell size so that its concentration is lower in larger cells to promote cell-cycle entry. Here, we find that a transcriptional control uncouples Whi5 synthesis from cell size and, screening for similar genes, identify histones as the major class of sub-scaling transcripts besides *WHI5*. Histone synthesis is thereby matched to genome content rather than cell size. Such sub-scaling proteins are challenged by asymmetric cell division because proteins are typically partitioned in proportion to new-born cell volume. To avoid this fate, Whi5 uses chromatin-binding to partition similar protein amounts to each new-born cell regardless of cell size. Finally, disrupting both Whi5 synthesis and chromatin-based partitioning compromises G1 size control. Thus, specific transcriptional and partitioning mechanisms determine protein sub-scaling to control cell size.

---

A striking feature of cell growth is that total protein and RNA amounts per cell increase approximately in proportion to cell volume as a cell grows (Fig. 1A) (Fraser and Nurse, 1978, 1979). To achieve this coordinated scaling of macromolecules with cell size, larger cells have higher transcription and protein synthesis rates (Creanor and Mitchison, 1982; Elliott, 1983; Elliott and McLaughlin, 1979; Elliott et al., 1979; Padovan-Merhar et al., 2015; Sun et al., 2020; Zhurinsky et al., 2010). This size-scaling is of general importance because it ensures macromolecule copy number is proportional to cell volume and therefore concentrations are kept constant as a cell grows (Fig. 1B) (Marguerat and Bahler, 2012; Neurohr et al., 2019). Nuclear volume also scales in proportion to cell volume meaning that nuclear concentrations are also expected to be constant (Jorgensen et al., 2007; Neumann and Nurse, 2007).

**Figure 1.**
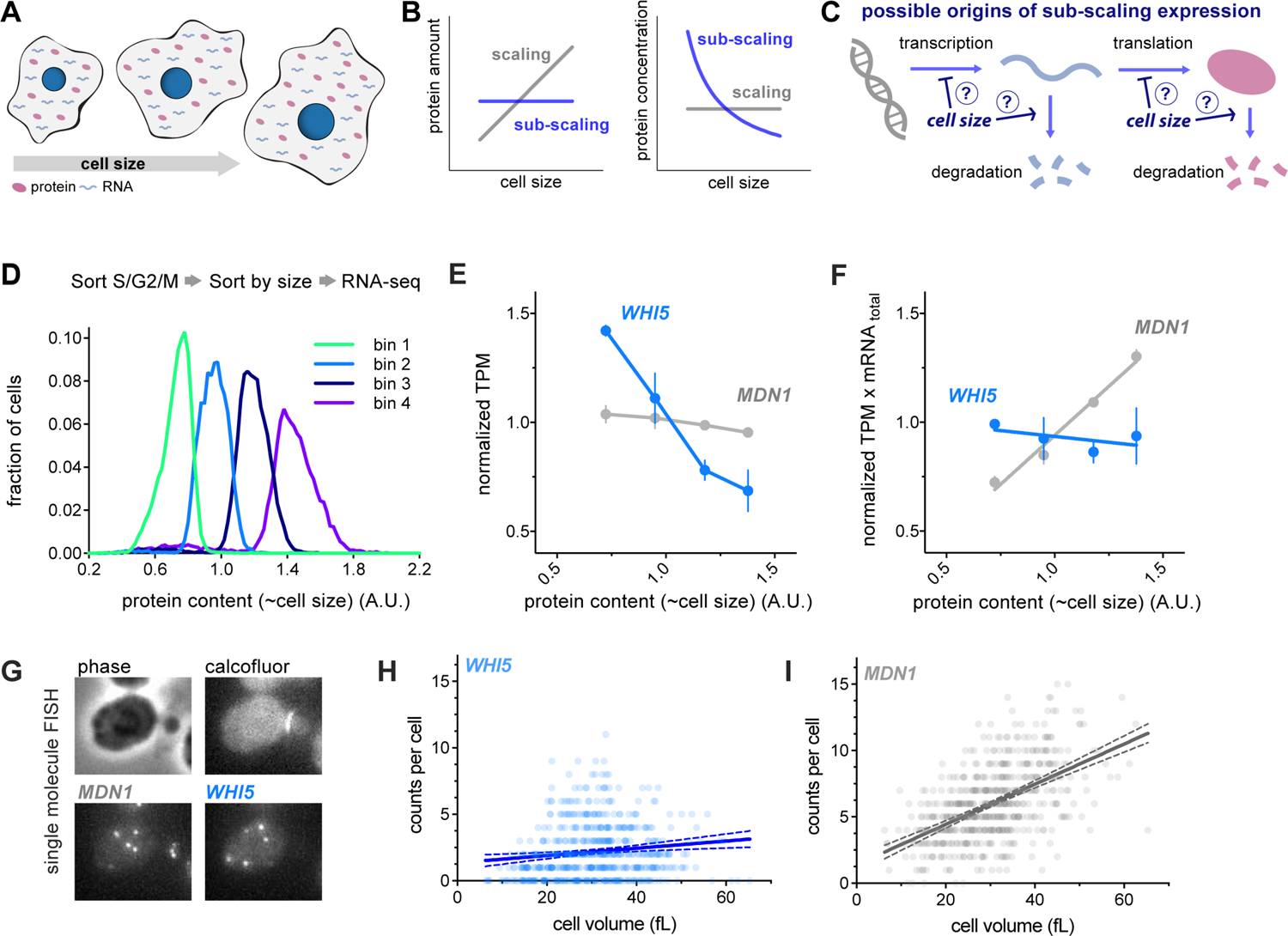
*WHI5* mRNA does not scale with cell size. (A-C) Schematics illustrating scaling and sub-scaling gene expression. (A) Total protein and RNA copy numbers per cell generally scale with to cell volume so that concentrations remain constant during growth. (B) However, some proteins sub-scale with size such that protein amounts are constant as a function of size and therefore protein concentrations decrease with cell size, which (C) could result from regulation at any step of gene expression. (D-F) Cells in S/G2/M were sorted into four bins based on the intensity of total protein dye. See Fig. S1 and Materials and Methods for details. (D) Histogram of total protein content per cell in each bin remeasured after sorting. (E) Normalized <u>T</u>ranscripts <u>P</u>er <u>M</u>illion (TPM / mean TPM) for *WHI5* and *MDN1* mRNA in cells of different sizes (total protein content). The mean (±range) of two biological replicates is plotted. Changes in TPM are proportional to changes in mRNA concentration. (F) Normalized TPM x total-mRNA for *WHI5* and *MDN1* mRNA in cells of different sizes (total protein content). Mean (±range) of two biological replicates is plotted. Changes in TPM x total mRNA are proportional to changes in mRNA amount. Relative total mRNA per cell was determined by the number of reads relative to those from a fixed number of *S. pombe* cells added to the sample. (G-I) single-molecule Fluorescence In Situ Hybridization (smFISH) analysis of *WHI5* and *MDN1* mRNA. (G) representative smFISH images. (H&I) mRNA counts per cell as a function of cell volume for *WHI5* and *MDN1* determined by smFISH, n=567 cells. Linear regression (solid line) and 95% confidence interval (dashed lines) are shown. Data are pooled from two biological replicates. The same data with replicates plotted independently are shown in Fig. S3A&B.

The importance of this biosynthetic size-scaling is underscored by experiments where cells are genetically manipulated to be excessively large. In these cases, protein and RNA synthesis can no longer keep pace with the expanding cell volume and the cytoplasm starts to dilute (Neurohr et al., 2019; Zhurinsky et al., 2010). This cytoplasmic dilution results in the failure of many key cellular processes including cell cycle progression and conditional gene expression programs (Neurohr et al., 2019; Zhurinsky et al., 2010). Importantly, this breakdown in biosynthesis only occurs in extremely large cells. To prevent themselves from becoming excessively large, proliferating cells coordinate cell division with cell growth so that cells born small compensate for their initially smaller size by growing more before entering the cell division cycle (Johnston et al., 1977; Turner et al., 2012).

The processes of cell size control and of biosynthetic scaling are deeply connected because one mechanism of size control relies on the differential scaling of cell cycle regulators with cell size. While it is generally assumed that most individual proteins exhibit the general size-scaling behavior and therefore remain at constant concentration, one notable exception is the cell cycle inhibitor Whi5 whose synthesis *sub-scales* with cell size (Schmoller et al., 2015) (Fig. 1B). Whi5 synthesis occurs during the S/G2/M stages of the cell cycle before Whi5 is translocated into the nucleus at the end of mitosis to inhibit the cell cycle SBF transcription factor in the following G1. Quantification of the synthesis rate of Whi5 in individual cells has revealed that Whi5 synthesis *sub-scales* with cell size so that an approximately constant amount of Whi5 is made in each cell cycle independent of cell size and global biosynthetic capacity (Schmoller et al., 2015). Whi5 concentration is then diluted in G1 as cells grow, reducing the inhibition of SBF activity and promoting cell-cycle entry in larger cells. Interestingly, examination of Whi5 synthesis in a variety of extracellular growth conditions showed that a similar number of molecules are made in all conditions. Whi5 synthesis is thus uncoupled from the cellular growth rate as well as cell size (Qu et al., 2019). In contrast, most other proteins’ synthesis is assumed to increase with cell size due in accordance with the global scaling trend. Thus, cell size determines the pattern of differential protein biosynthesis, which promotes division in larger cells to, in turn, control cell size.

This idea of cell cycle regulators differentially scaling during G1 has been expanded on by recent work in budding yeast examining cells arrested in G1 for increasing amounts of time. This revealed that as a G1 arrest is prolonged, some cell cycle activators increase in concentration while some cell cycle inhibitors decreased in concentration to promote the cell cycle entry of larger cells (Chen et al., 2020). Such a size-dependent concentration increase was first observed in the fission yeast *S. pombe* for the cell cycle activator *cdc25* (Keifenheim et al., 2017).

Despite this progress, the underlying molecular mechanisms determining the relationship between cell size and the expression of individual proteins remain largely unknown. It is both unclear what mechanisms scales most biosynthesis with cell size and also what additional mechanisms uncouple the synthesis of sub-scaling proteins such as Whi5 from the general trend. It also remains unknow how pervasive sub-scaling behavior is and which other categories of proteins sub-scale to differentially coordinate other aspects of cell biology with cell size.

To address these questions at the heart of how protein synthesis scales with cell size, we have used multiple orthogonal high-throughput and single-cell approaches. We identified a transcriptional control mechanism uncoupling Whi5 synthesis from cell size. Besides *WHI5,* we identified histones as the major class of sub-scaling gene products by analyzing the transcriptome of differently sized cells progressing through the cell cycle. For stable proteins such as Whi5, we show how asymmetry in cell division presents a challenge to their sub-scaling expression. This is because the default manner in which proteins are partitioned is in proportion to the volume of the new-born cell which would result in the smaller daughter cell inheriting proportionally less protein – effectively undoing the sub-scaling synthesis of the preceding cell cycle. To avoid this fate, Whi5 uses chromatin-binding to segregate a similar number of protein molecules to each new-born cell regardless of their size. Finally, we disrupted both Whi5 synthesis sub-scaling and Whi5’s chromatin-based partitioning and show that together these mechanisms are required for G1 size control in budding yeast.

## *WHI5* mRNA does not scale with cell size

First, we set out to determine at what stage of gene expression Whi5 sub-scaling originates. In principle, any step of gene expression could be regulated in a manner that results in sub-scaling protein levels (Fig. 1C). Importantly, for Whi5 this is not achieved through negative feedback on gene product levels because multiple copies of the *WHI5* gene result in a proportional increase in the number of proteins made per cell cycle (Qu et al., 2019; Schmoller et al., 2015). To determine whether Whi5’s sub-scaling behavior originates at the protein or transcript level we isolated cells of different sizes by FACS using total cellular protein content as a proxy for cell size. To do this we stained cells with an amine reactive NHS ester dye which binds bulk protein, sorted cells into four bins based on dye intensity and performed RNA-seq on each bin (Fig. 1D & S1A). The protein dye intensity in each bin was well correlated with total mRNA content and cell volume, confirming the protein dye as a good general proxy for cell size (Fig. S1B-E). Total mRNA was quantified by adding a *S. pombe* spike-in before RNA extraction and then determining the ratio of cerevisiae to *S. pombe* reads, and cell volume was measured by coulter counter after sorting. *WHI5* mRNA transcripts per million (TPM) decreased as cell size increased, which implies that the *WHI5* mRNA concentration is lower in larger cells. In contrast, *MDN1* mRNA TPM, as representative of scaling gene expression, was constant (Fig 1E).

We then normalized the TPM value to the total mRNA amount per cell, determined relative to the *S. pombe* spike-in, to get an estimate of the relative transcript amount per cell. When normalized to total mRNA, *WHI5* mRNA amount per cell was constant as a function of size (Fig. 1F). To corroborate this finding, we performed single-molecule FISH in individual cells while also measuring the size of each individual cell (Fig 1G). Consistent with the RNA-seq data, the number of *WHI5* transcripts per cell did not increase with cell size whereas the *MDN1* counts did (Fig. 1H-I & S3A-B).

## Cell cycle analysis of *WHI5* sub-scaling

Next, we sought to test if the apparent sub-scaling behavior of *WHI5* mRNA is simply a consequence of the cell cycle rather than cell size *per se.* This is a possibility because *WHI5* mRNA is a cell cycle regulated transcript that peaks in S phase (Pramila et al., 2006) (Fig. S3D) and cells later in the cell cycle will on average tend to be larger. To control for this possibility, we isolated cells in early G1 by centrifugal elutriation and arrested them in G1 for increasing amounts of time to generate populations of cells of increasing sizes. Cells were then released from the G1 arrest resulting in cultures of cells synchronously traversing the entire cell cycle but at different sizes, which we then sampled for RNA-seq analysis (Fig. 2A-B & S2). *WHI5* TPM are indeed lower in larger cells as *WHI5* expression peaks in S phase (Fig. 2C). The total amount of *WHI5* expression across the entire cell cycle can then be estimated as the area under the curve, which again shows that *WHI5* mRNA concentrations decrease as cell size increases (Fig. 2D).

**Figure 2.**
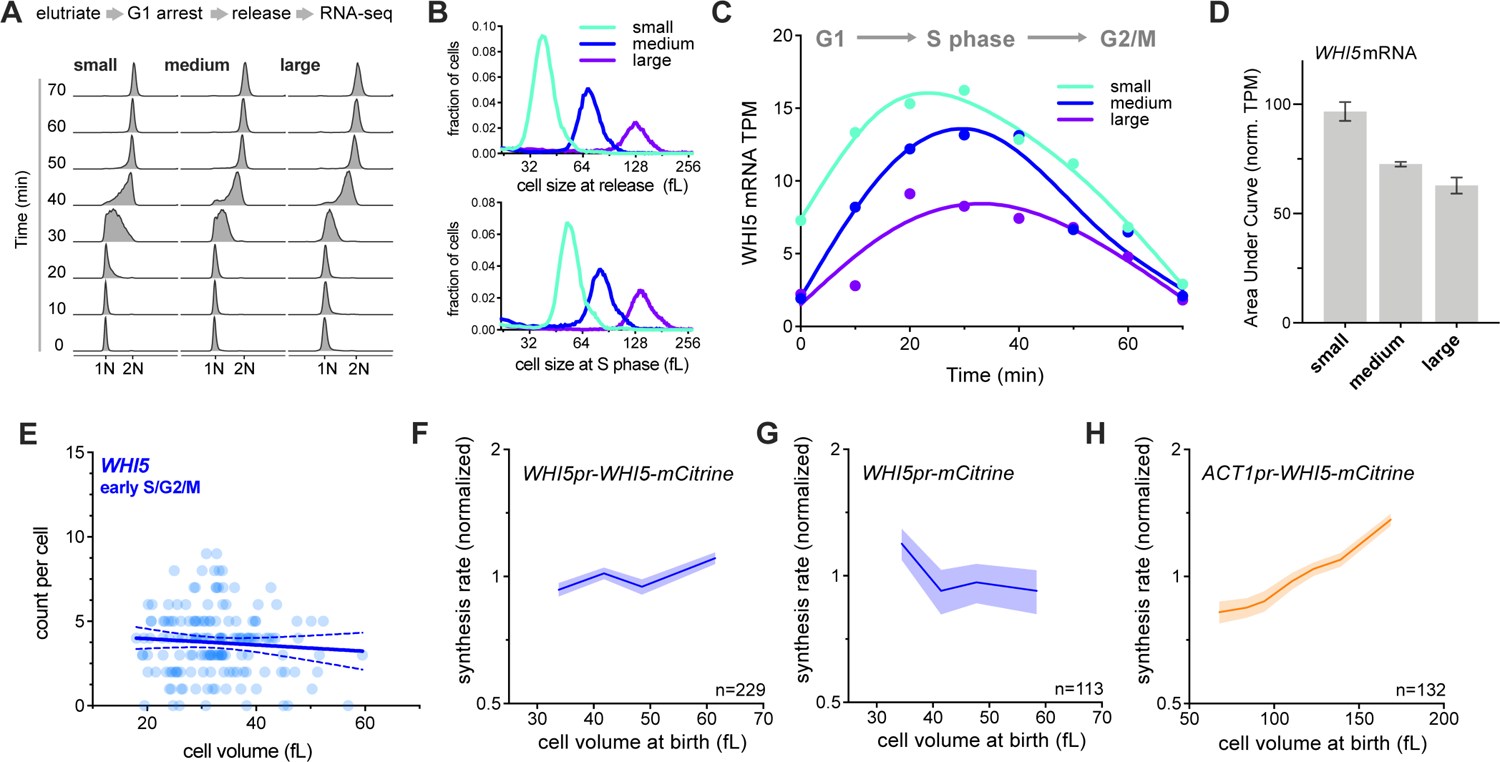
*WHI5* sub-scaling occurs across the cell cycle and is encoded in the *WHI5* promoter. (A-D) G1 cells of different sizes (small, medium, and large) were arrested for increasing amounts of time in G1 using a temperature sensitive *cdc28-13* allele at 37°C. Cells were then released from G1 to progress synchronously through a full cell cycle and analyzed by RNA-seq. See Fig. S2 for details. (A) DNA content analysis determined by flow cytometry. (B) Size distributions at point of release from G1 arrest (top panel) and at mid S-phase (bottom panel, corresponds to the 40-minute time point). (C) *WHI5* mRNA TPM and (D) the <u>A</u>rea <u>U</u>nder the <u>C</u>urve (AUC) of mean normalized *WHI5* mRNA TPM for small, medium large cells synchronously progressing through the cell cycle. The AUC mean (± range) of two biological replicates is plotted. (E) mRNA counts per cell for *WHI5* as a function of cell size in early S/G2/M cells determined by smFISH; n=156 cells. Early S/G2/M cells were defined as budded cells with a small (≤ 0.2) bud-to-mother volume ratio. Linear regression (solid line) and 95% confidence interval (dashed lines) are shown. Data are pooled from two biological replicates. The same data with replicates plotted independently, including data for *MDN1*, are shown in Fig. S3E&F. (F-H) Protein synthesis rates normalized to the mean as a function of cell volume at budding were determined by time-lapse fluorescence microscopy measuring Whi5-mCitrine expressed from (F) the endogenous *WHI5* promoter or (H) the *ACT1* promoter, and (G) mCitrine expressed alone from the *WHI5* promoter. Synthesis rates were determined as in Schmoller et al., 2015 for single cells using linear fits of total protein traces for the period between bud emergence and cytokinesis (S/G2/M). Data are binned according to cell volume at budding and the mean (±SEM) of each bin is plotted. Un-binned single-cell values from the same data are plotted in Fig. S3G-I.

Consistent with this, if we restrict our smFISH analysis to only those cells in early S/G2/M, when *WHI5* expression peaks, the number of *WHI5* transcripts is still uncorrelated with cell size (Fig. 2E & S3E-F). Taken together, this group of experiments suggests that *WHI5* transcript levels are responsible for the sub-scaling expression of Whi5 protein.

## *WHI5* sub-scaling is encoded in its promoter

Next, we sought to test whether Whi5 sub-scaling is encoded in its promoter. If this were the case, then the *WHI5* promoter should be both necessary and sufficient for sub-scaling protein expression. To test if the *WHI5* promoter is sufficient, we compared the size-dependency of Whi5-mCitrine protein expressed from its endogenous promoter with that of a reporter mCitrine also expressed from the *WHI5* promoter. Both Whi5-mCitrine and the mCitrine reporter are synthesized in a sub-scaling manner indicating that the WHI5 promoter is sufficient for WHI5 sub-scaling synthesis (Fig. 2F-G). In contrast, when we expressed Whi5-mCitrine from a scaling promoter (*ACT1pr*) its synthesis rate increases with cell size (Fig. 2H). Together, these experiments suggest that *WHI5* is transcribed in a sub-scaling manner and that the *WHI5* promoter is both necessary and sufficient for the sub-scaling synthesis pattern of Whi5.

## Histones are a rare class of sub-scaling genes

Having shown that sub-scaling expression of *WHI5* is due to a transcriptional mechanism, we sought to determine which other cellular processes are similarly uncoupled from cell size. To do this, we analyzed our RNA-seq experiments of different sized cells. We found 15 transcripts that behaved similarly to *WHI5* in both the size-sort and the elutriation arrest-release experiment (Figure S4A). These genes are enriched for GO terms related to chromatin and revealed histones as the major class of sub-scaling genes, as 9 of the 15 identified genes encode histones (Fig. 3A). Histone mRNAs TPMs are clearly lower in the larger sorted cells than the smaller ones (Fig. 3B & S4B). As for *WHI5,* this is not a consequence of their cell cycle regulated expression because the same trend was observed in the timecourse experiments where cells of different sizes synchronously progressing through the entire cell cycle (Fig. 3C-D & S4C).

**Figure 3.**
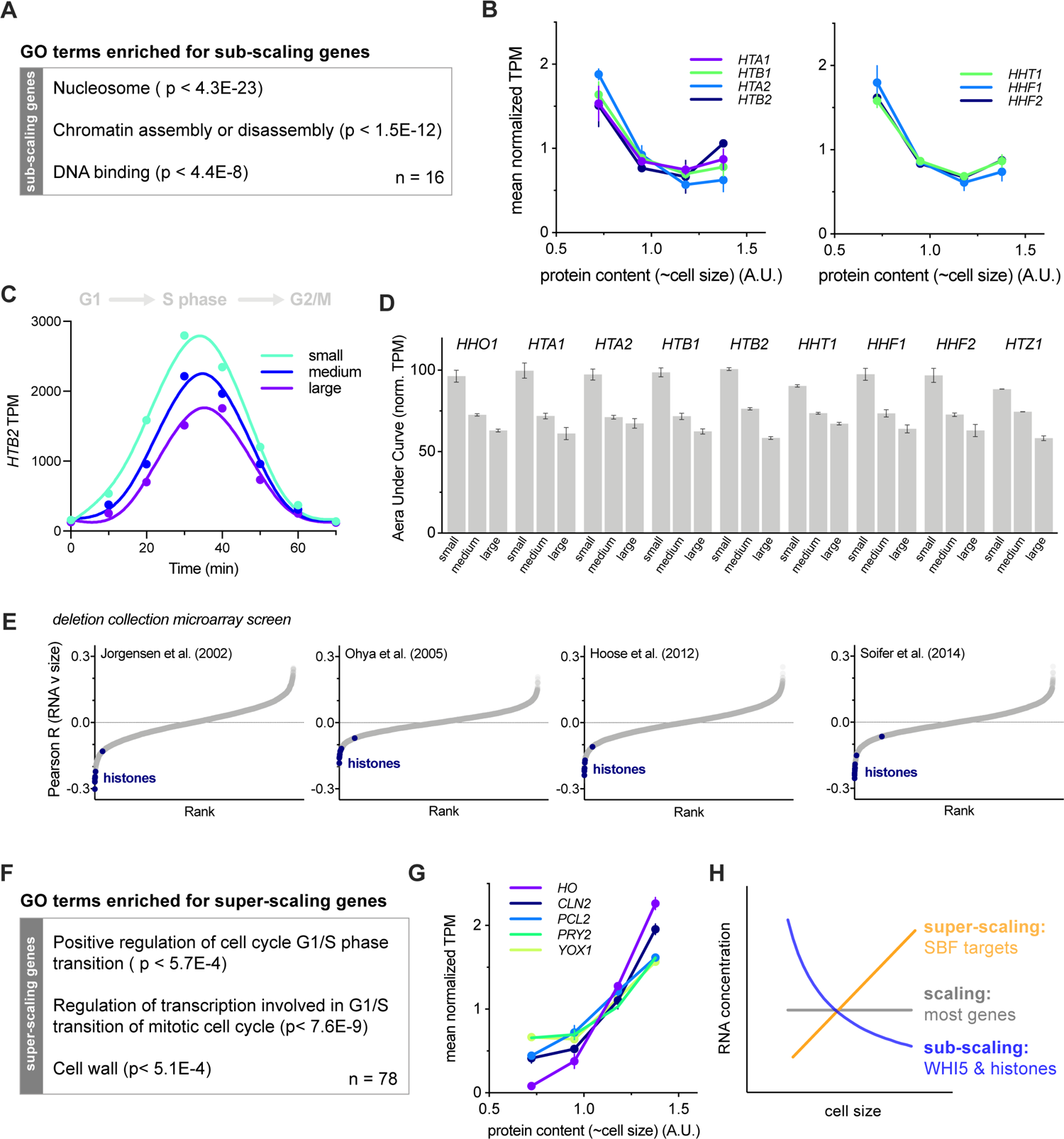
Histones are a rare class of sub-scaling genes. (A) Gene ontology terms enriched in sub-scaling genes. 9 of the 16 sub-scaling genes encode histones and one is *WHI5*. See Fig. S4A and Materials and Methods for classification details. (B) Normalized TPM (TPM / mean TPM) for sub-scaling histone mRNAs in cells of different sizes (total protein content). The mean (±range) of two biological replicates is plotted. Changes in TPM are proportional to changes in mRNA concentration. See Fig. S1 for experimental details. (C) *HTB2* mRNA TPM for small, medium, and large cells synchronously progressing through the cell cycle as in Fig.2C&D. See Fig. S2 for experimental details. (D) The Area Under the Curve (AUC) of mean normalized sub-scaling histone mRNA TPM of small, medium, and large cells synchronously progressing through the cell cycle. The AUC mean (±range) of two biological replicates is plotted. (E) Pearson correlation coefficient, R for the correlation between Histone mRNA levels, relative to wild-type, in 1,484 gene deletion strains (Kemmeren et al., 2014; O’Duibhir et al., 2014) and the cell size of the respective gene deletions for four different data sets of size measurements (Hoose et al., 2012; Jorgensen et al., 2002; Ohya et al., 2005; Soifer and Barkai, 2014). Each point represents an individual transcript. Histone mRNAs are shown in blue. The individual regression fits for the histone transcript levels with cell size determined by Jorgensen et al. are shown in Fig. S4D. (F) Gene ontology terms enriched in super-scaling genes. See Fig. S6A and Materials and Methods for classification details. (G) Normalized TPM (TPM / mean TPM) for example super-scaling mRNAs, specifically those known as targets of the SBF transcription factor, in cells of different sizes (total protein content). The mean (±range) of two biological replicates is plotted. Changes in TPM are proportional to changes in mRNA concentration. See Fig. S1 for experimental details. (H) schematics illustrating the scaling, sub-scaling and super-scaling trends of gene expression, representative of most genes, WHI5 and histones, and a subset of SBF targets respectively.

We further confirmed the sub-scaling expression of histone transcripts by examining microarray data of 1,484 strains each containing a single gene deletion (Kemmeren et al., 2014; O’Duibhir et al., 2014). We compared the level of a given transcript in each deletion strain with the cell size of the same deletion strain and then calculated the Pearson R coefficient for the correlation between transcript levels and cell size across all 1,484 deletion strains. We repeated this using four different cell size datasets acquired as part of independent genome-wide screens utilizing multiple different methodologies for measuring cell size (Hoose et al., 2012; Jorgensen et al., 2002; Ohya et al., 2005; Soifer and Barkai, 2014). This revealed a clear negative correlation between histone mRNA levels and cell size (Fig. 3E & S4D), meaning that histone mRNA concentrations are lower relative to the rest of the transcriptome in deletion strains with a larger cell size. Indeed, histones populate the most extreme negative end of the spectrum of transcripts in all four datasets (Fig. 3E). This contrast to typical transcripts, such as those encoding RNA polymerase II subunits (Fig. S5).

We next sought to identify *super-scaling* genes whose mRNA concentrations increase in larger cells (Fig. S6A). To do this, we again analyzed how gene expression through the cell cycle changes as a function of cell size (Fig. S1 & S2). This identified a number cell cycle regulated transcripts including SBF regulated genes such as *CLN2* that super-scale (Fig. 3F-H & S6B-C). That these cell cycle regulated genes super-scale whereas histones and *WHI5* sub-scale, despite both sets peaking in expression at a similar time, demonstrates that these differential scaling properties are not due to a conflation of cell cycle progression with cell size. Consistent with this conclusion, many other cell cycle regulated transcripts, including the B-type cyclins, do not sub-scale (S6D-E).

## Histone protein synthesis is similarly uncoupled from cell size

The sub-scaling expression of histone mRNAs suggests that histone protein expression is coordinated with genome content rather than cell size and predicts that histone protein synthesis should also not scale proportionally with cell size. To examine this, we first analyzed two published datasets of flow cytometry measurements across the collection of strains in which each individual open reading frame was fused to GFP (Parts et al., 2014). We compared the relationship between GFP fluorescence (protein amount) and side scatter (SSC-A, cell size) (Fig. 4A). This revealed that histone protein amounts show weaker dependence on cell size than the average protein in the proteome, *i.e.*, the slope between cell size (SSC-A) and GFP intensity is smaller (Fig. 4B-C). To confirm that histone protein synthesis does indeed sub-scale, we quantified the amount of histones synthesized across the cell cycle in single cells using time-lapse fluorescence microscopy and compared it with the increase in cell volume during the same period. While the protein synthesis of the RNA polymerase II subunit Rpb3 scales with cell size, the protein synthesis of histones Hta2, Htb2 and Htz1 clearly sub-scale with size (Fig. 4D-E). Taken together, these experiments identified histones as a rare class of sub-scaling genes whose transcription and protein synthesis are uncoupled from cell size. In this way, histone production can be matched with genome-content rather than cellular growth.

**Figure 4.**
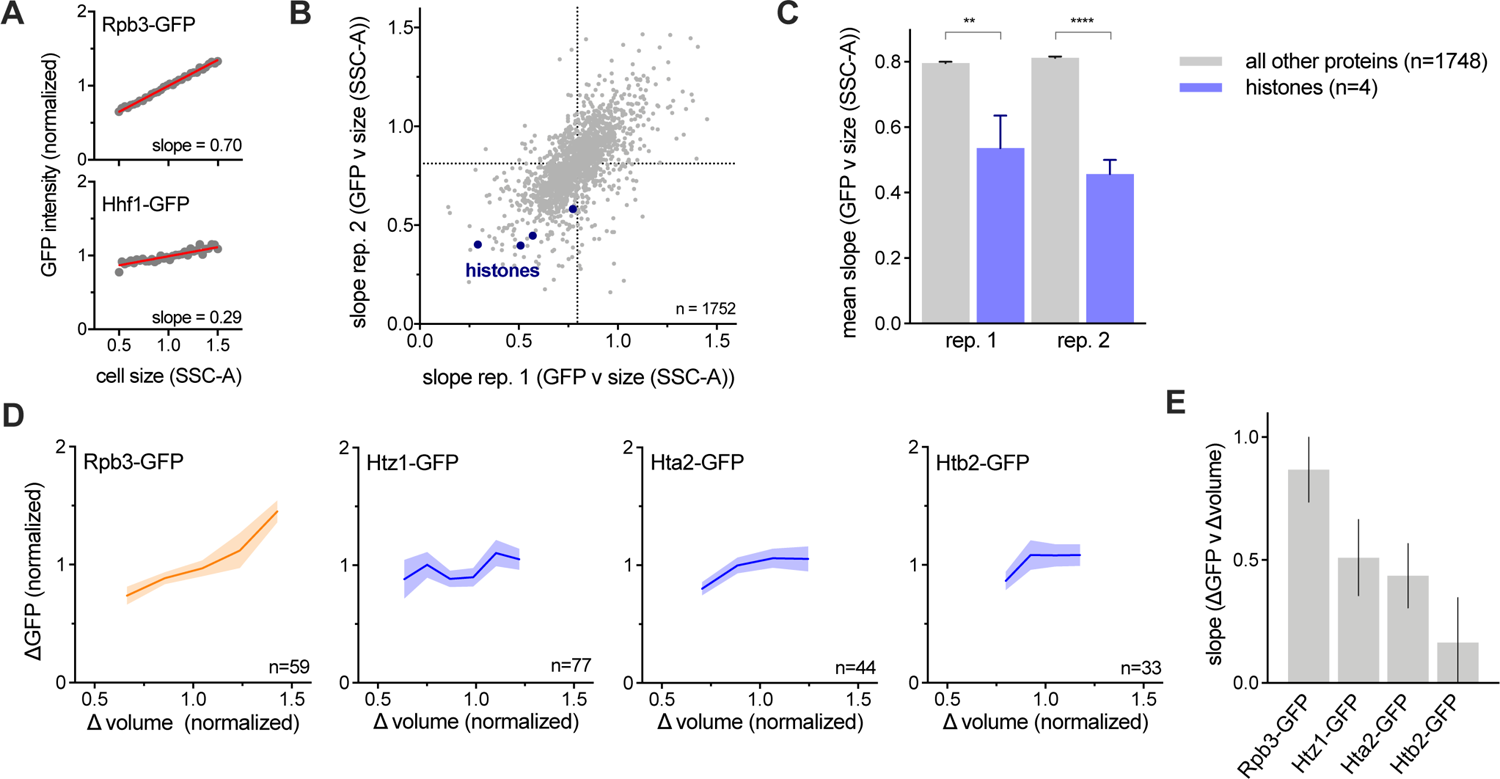
Histone protein synthesis does not scale with cell size. (A-C) Analysis of size-dependent expression in the genome-wide collection of GFP fusion strains measured by flow-cytometry (Parts et al., 2014). Side scatter (SCC-A) was used as a proxy for cell size. The slope of the linear fit between cell size (SCC-A) and GFP intensity in budded cells was used to estimate the degree of size-dependence for each protein. See Materials and Methods for details. (A) Plot of example protein-GFP levels (intensities normalized to the mean intensity) against cell size (SCC-A normalized to the mean SCC-A) in budded cells. Grey dots denote bin means. Red lines show the linear regression to the un-binned data. (B) Slope values (for the linear regression between GFP and cell size in budded cells) of 1752 proteins analyzed in two replicates. Slopes closer to 0 correspond to stronger sub-scaling behavior. Histone proteins are shown in blue. (C) Average slope values (for the linear regression between GFP and cell size in budded cells) for histones (blue) and all other proteins (grey). Four histone proteins were present in the 1752 proteins analyzed. Histone proteins have significantly smaller slopes than the average protein (** p=0.0014; **** p <0.0001). (D) The amount of Rpb3-GFP (RNA polymerase II subunit) and three histones (Hta2-GFP, Htb2-GFP and Htz1-GFP) synthesized (ΔGFP normalized to its mean) between birth and division as a function of the amount of growth (Δvolume normalized to its mean), determined by single cell time-lapse fluorescence microscopy. Data are binned according to Δvolume and the bin means (±SEM) are plotted. (E) Slope of the linear fits to single cell values of ΔGFP against Δvolume. Error bars show the standard error of the slope.

## Inheritance of sub-scaling protein levels requires chromatin-based partitioning

Both Whi5 and histones are stable proteins synthesized in a sub-scaling manner during S/G2/M of the cell cycle, meaning that their amounts in G1 are determined by inheritance from previous cell cycles. For typical proteins, which are partitioned along with the cytoplasm or nucleoplasm, concentrations are expected to be similar in the mother and daughter cells following division, as is observed for a freely diffusing mCitrine (Fig. 5A&B). Thus, the asymmetric division of budding yeast poses a problem for maintaining the protein-level sub-scaling of Whi5 and histones because smaller daughter cells would inherit fewer proteins if they were partitioned in proportion to cell volume. Instead, to maintain size-independent amounts, a mechanism partitioning equal amounts to the daughter and mother cells is required. This is indeed the case for Whi5, which is not partitioned evenly by volume as seen by the increased mother-to-bud concentration ratio at cytokinesis (Fig. 5B). Thus, differently sized G1 cells inherit a more similar amount of Whi5 during cell division than would be expected for any typical protein partitioned by volume.

**Figure 5.**
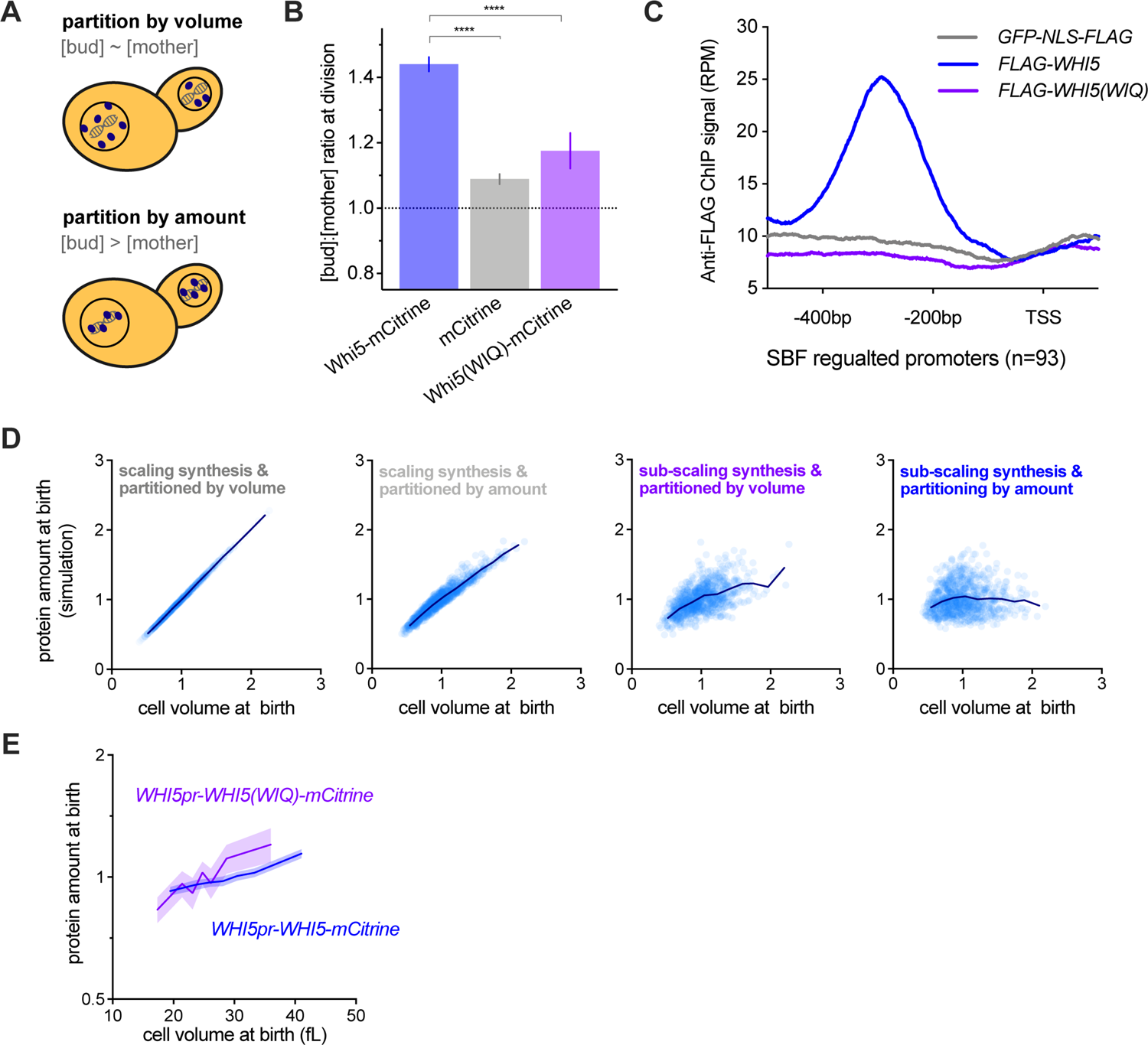
Maintenance of Whi5 sub-scaling requires chromatin-based partitioning during asymmetric division. (A) Schematic illustrating the different regimes of protein partitioning at cell division which can be quantified by comparing the mother-to-bud protein concentration ratio at cytokinesis. A ratio ∼ 1 is expected for proteins that are partitioned in proportion to volume. A ratio > 1 is expected for proteins that are partitioned in equal protein amounts (*i.e.*, independent of the mother and bud volume) (B) The bud-to-mother concentration ratios for Whi5-mCitrine, free mCitrine, and Whi5(WIQ)-mCitrine at cytokinesis. Whi5(WIQ)-mCitrine has reduced recruitment to DNA because it does not bind the SBF transcription factor (Travesa et al., 2013) (Fig. 5C & S7A). **** p < 0.0001. (C) anti-Flag ChIP-seq experiments were performed to compare Whi5, Whi5(WIQ) and a GFP-NLS ChIP control. Average RPM metagene plot upstream of all SBF regulated promoters (as defined by (Ferrezuelo et al., 2010)) is shown. ChIP signal around individual SBF binding sites, including additional replicates and controls, is shown in Fig S7A. (D) Simulation protein amounts at birth as a function of daughter volume at birth. Four different conditions were simulated where protein expression was either in proportion to cell size (scaling) or independent of cell size (sub-scaling), and protein partitioning is either by amount or in proportion to cell volume. Individual simulated cells (light blue) as well as bin means (dark blue) are plotted. Protein concentrations from the same simulation are shown in Fig. S7B. (E) Protein amount at birth (normalized to the mean) as a function of cell volume at birth for *WHI5pr-WHI5-mCitrine* and *WHI5pr-WHI5(WIQ)-mCitrine* cells. Data are binned according to cell size at birth and the bin means (±SEM) are plotted. Un-binned single-cell values of the same data are plotted in Fig. S7D.

To obtain a quantitative understanding of the impact of amount- and volume-based partitioning modalities on the amounts of inherited protein, we employed a full cell-cycle model that simulates growth and division of a population of budding yeast cells. This model was parameterized by single-cell microscopy measurements and therefore accounts for cell-to-cell variability and the size-dependence of cell cycle progression (Chandler-Brown et al., 2017). To this model, we added a protein synthesized either at a rate proportional to cell size (scaling) or independent of cell size (sub-scaling). At division, these proteins were then either partitioned in proportion to cell-volume or in the manner empirically determined for Whi5, where most but not all, of the protein is partitioned by amount (Fig. 5D & S7B). Our simulations show that the amount of Whi5 inherited in G1 should scale significantly more with cell size if it were partitioned according to cell volume rather than by amounts. This is, in part, because bud size varies significantly even for mothers with the same volume. Together, this suggests that both partitioning by amount and sub-scaling synthesis should be required to ensure Whi5 protein sub-scaling in G1.

We hypothesized that the amount-based partitioning of Whi5 could be achieved by utilizing the equal partitioning of the genome during cell division. This possibility was suggested by the fact that Whi5 binds the DNA-bound SBF transcription factor complex following its dephosphorylation at mitotic exit. To test this, we analyzed the partitioning of a Whi5 mutant, Whi5(WIQ), that does not bind SBF and is not recruited to the SBF binding sites in the *CLN2* or *SVS1* promoters (Travesa et al., 2013). First, we confirmed that the WIQ mutation reduces Whi5 binding at SBF bound DNA elements across the genome by ChIP-seq (Fig. 5C & S7A). Next, we analyzed single cells expressing Whi5(WIQ)-mCitrine, which revealed Whi5(WIQ)-mCitrine has a significantly lower bud-to-mother concentration ratio at division than wild-type Whi5. This supports our model that partitioning by amount is indeed mediated by chromatin binding (Fig. 5A). Crucially, Whi5(WIQ) amounts at birth are higher in larger cells and lower in smaller cells when compared to wild-type Whi5, demonstrating that when chromatin-binding based partitioning of Whi5 is disrupted, its sub-scaling in G1 is also disrupted (Fig. 5E). Thus, our data support a model where Whi5 binding to chromatin results in its volume-independent partitioning into the mother and bud at division to ensure daughter cells inherit approximately the same amount of Whi5 regardless of their size.

## G1 size control requires Whi5 sub-scaling

After establishing that Whi5 expression relies on a combination of sub-scaling transcription and chromatin-based partitioning, we proceeded to disable both mechanisms to test the function of Whi5 sub-scaling in G1 size control by generating cells with constant Whi5 concentrations (Fig. 6). We used the *bck2Δ* background because *BCK2* and *WHI5* may be involved in parallel size control pathways (Schmoller et al., 2015) and we are here focusing on the *BCK2*-independent branch of G1 size control. We have also focused our analysis specifically on daughter cells growing in a poor non-fermentable carbon source (1% ethanol + 2% glycerol) because under these conditions G1 size control is most pronounced, in part due to the smaller and more variable size of new born daughters.

**Figure 6.**
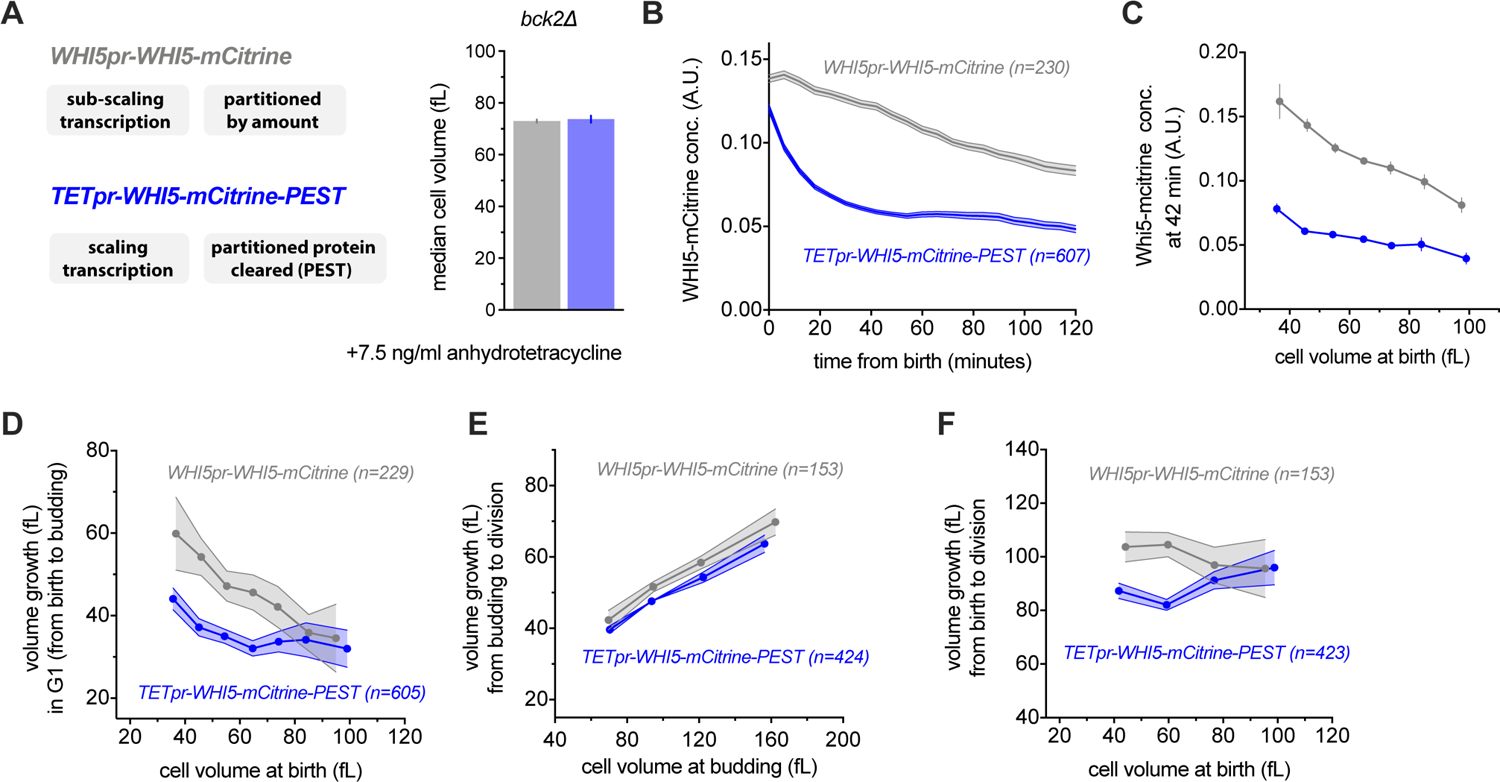
Disruption of Whi5 sub-scaling compromises G1 size control. (A) Median cell volume (average of two independent measurements), measured by Coulter counter of cells expressing *WHI5pr-WHI5-mCitrine* (grey) or *TETpr-WHI5-mCitrine-PEST (*blue) in the presence of 7.5ng/ml anhydrotetracycline to induce expression from the size-scaling *TET* promoter. The fused *PEST* domain destabilizes Whi5 to eliminate Whi5 synthesized in the preceding cell cycle from new-born G1 cells. (B-F) Single cell time-lapse microscopy was performed on *bck2Δ* strains expressing either *WHI5pr-WHI5-mCitrine* or *TETpr-WHI5-mCitrine-PEST.* D-F show binned data where no more than 2 cells do not fit within the cell volume bin limits. (A) (B) Mean Whi5 concentration (±SEM) as a function of time from birth in all cells that completed G1. (B) Mean Whi5 concentration (±SEM) at 42 minutes after birth as a function of cell volume at birth in all cells that completed G1. (C) Cell growth in G1 as a function of cell volume at birth for cells that completed G1. Data are binned according to cell volume at birth and the bin means (±SEM) are plotted. Un-binned single-cell values of the same data are plotted in Fig. S8B&C. (D) Cell volume growth between budding and cell division (S/G2/M) as a function of cell volume at budding. Data are binned according to cell volume at budding for cells at completed the cell cycle and the bin means (±SEM) are plotted. (E) Cell volume growth between birth and cell division as a function of cell volume at birth. Data are binned according to cell volume at birth for cells with a completed cell cycle and the bin means (±SEM) are plotted.

The memory of Whi5 partitioning throughout G1 phase relies on it being a stable protein. We therefore destabilized Whi5 by fusing Whi5-mCitrine with the Cln2 PEST degron sequence (Mateus and Avery, 2000). This allowed us to bypass size-independent partitioning because any Whi5 inherited from the previous cell cycle should then be degraded during early G1. We then expressed the Whi5-mCitrine-PEST fusion from a synthetic TET promoter that is conditionally activated by anhydrotetracycline in a dose-dependent manner, and whose expression scales with cell size like most genes (Azizoğlu et al., 2020). We grew cells in 7.5 ng/ml anhydrotetracycline because at this concentration the average cell size was similar to that of *bck2Δ* cells expressing *WHI5-mCitrine* expressed from its own promoter (*WHI5pr-WHI5-mCitrine*) (Fig. 6A). Thus, the *TETpr-WHI5-mCitrine-PEST* strain allows us to generate G1 cells where Whi5 amounts should no longer sub-scale with size. Instead, Whi5 concentrations in new-born daughter cells start high due to the asymmetrically partitioning, but it is then degraded in the first ∼40 minutes of G1 after which a steady-state Whi5 concentration is established (Fig. 6B-C). This is in contrast to *WHI5-mCitrine* expressed from its own promoter, which is steadily diluted as cells grow during G1 (Fig. 6B).

When we examined size control in *WHI5pr-WHI5-mCitrine* cells, a clear anticorrelation between cell volume at birth and volume growth in G1 was observed, as expected (Figs. 6D & S8A). In contrast, this anticorrelation between cell volume at birth and volume growth in G1 was significantly weaker in *TETpr-WHI5-mCitrine-PEST* cells, showing that Whi5 sub-scaling and dilution is important for cell size control in G1 (Figs. 6D & S8B). We did not observe any significant perturbation to the relationship between size at budding and growth during S/G2/M (Fig. 6E). As previously reported, for wild-type cells the two independent phases of G1 and S/G2/M combine to form an apparent adder, where an approximately fixed absolute amount of volume growth occurs during each entire cell cycle (Chandler-Brown et al., 2017). In contrast, *TETpr-WHI5-mCitrine-PEST* cells display a positive correlation between size at birth and total cell cycle volume growth (Fig. 6F).

The fact that the anticorrelation between birth size and G1 growth is not completely lost but only reduced (Figs. 6D & S8A-C) is consistent with the notion that other pathways feed into the G1/S transition to couple growth and cell cycle entry (Chen et al., 2020). Nevertheless, our data clearly indicates that Whi5 dilution and sub-scaling contribute significantly to G1 cell size control in budding yeast.

## Discussion

In conclusion, while most proteins are synthesized proportionally to cell size, a handful are not and instead sub-scale with cell size (Fig. 1A&B). Such sub-scaling is already apparent in the mRNA amounts for both histones and *WHI5* across the cell cycle. Our *WHI5* promoter-swap experiments indicate that specific promoter elements are at least partly responsible for sub-scaling synthesis. In addition to this transcriptional control of synthesis, sub-scaling proteins also need a dedicated mechanism to ensure that equal protein amounts are inherited by differently sized cells during cell division. We discovered that Whi5 uses chromatin-binding as a mechanism to segregate approximately equal numbers of molecules to each new-born cell and thereby ensure protein synthesized in the preceding cell cycle is inherited by new-born cells independently of their size (Fig. 7).

**Figure 7.**
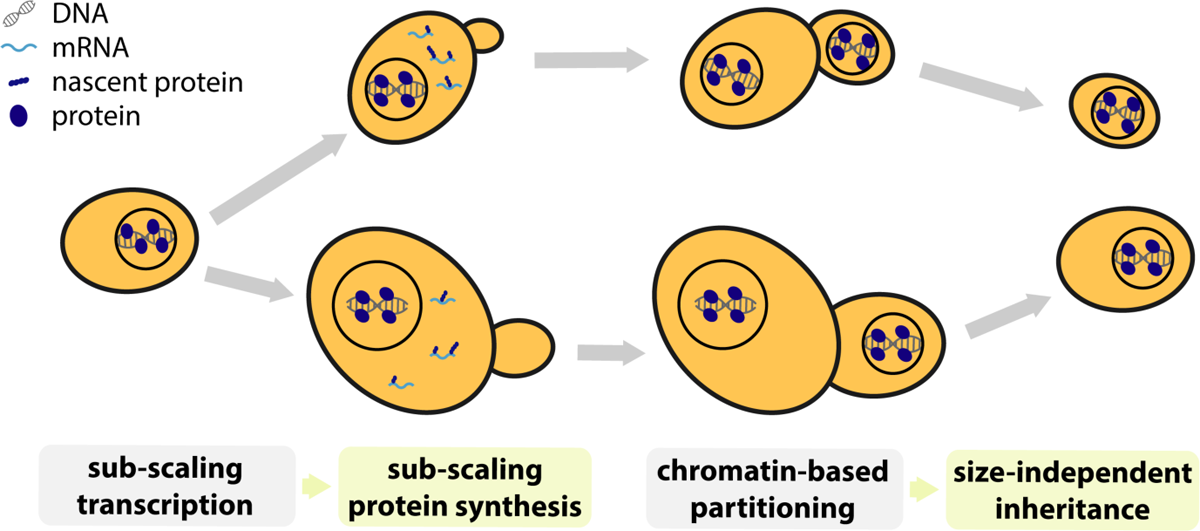
Summary schematic. A small class of genes including the cell cycle inhibitor *WHI5* and histones are transcribed in a sub-scaling manner, resulting in sub-scaling protein synthesis during the cell cycle. Sub-scaling proteins must also be partitioned during cell division independent of daughter cell size, which is achieved through chromatin-based partitioning.

### Histone genes dominate the sub-scaling gene class

Through a combination of multiple transcriptomic and proteomic screens we identified histones as the major class of sub-scaling genes in addition to *WHI5*. Sub-scaling of histone synthesis maintains a stoichiometric relationship between the amount of histones and the genome without engaging wasteful feedback mechanisms, which are known to operate when histone expression is artificially perturbed (Cross and Smith, 1988; Gunjan and Verreault, 2003; Moran et al., 1990; Norris and Osley, 1987). In this way, the amount of histone synthesis in each cell cycle better reflects the binary increase in genome content rather than variations in cell size. We speculate that this helps prevent unwanted variations in chromatin structure or accessibility in different sized cells.

### Whi5 sub-scaling is one of multiple inputs for G1 size control

In the case of *WHI5*, the function of sub-scaling synthesis is to control cell size. Whi5 functions in early G1 of the cell cycle to inhibit the SBF transcription factor and thereby delay cell cycle progression (Costanzo et al., 2004; de Bruin et al., 2004). Whi5 is inactivated by phosphorylation by cyclin-Cdk complexes that also drive its exclusion from the nucleus at *Start*, the point of commitment to cell division (Doncic et al., 2011). Importantly, the exclusion of Whi5 from the nucleus at *Start* marks the end of the most size-dependent part of the cell division cycle (Di Talia et al., 2007).

The sub-scaling of Whi5 allows its concentration to reflect and control cell size. That all cells are born with similar amounts of Whi5 protein, which is then diluted in G1, means that the Whi5 concentration is a readout of current cell size. Since Whi5 is a cell cycle inhibitor, its higher concentration in smaller cells delays their *Start* transition so that they have more time to grow in G1. Conversely, larger cells have lower concentrations of Whi5 and therefore more rapidly enter the cell division cycle. This importance of Whi5 sub-scaling for cell size control in G1 is demonstrated by our experiments where we synthetically disable Whi5 sub-scaling and observe a weakened G1 size control (Fig. 6). Crucially, to remove Whi5 sub-scaling, we had to bypass *both* the sub-scaling transcription and the chromatin-based partitioning mechanisms. Our data indicate that while Whi5 sub-scaling and dilution contribute significantly to budding yeast G1 cell size control, they do not remove it completely. This is consistent with the presence of additional size control mechanisms (Chen et al., 2020). We note that the experiments we present here are related to, but differ from a previous examination of the effect of *GAL1pr-WHI5* expression utilizing a stable Whi5 protein in a *BCK2+* background where the chromatin-based partitioning of Whi5 has not been removed (Barber et al., 2020). Taken together, theirs and our results are consistent with both the transcriptional and partitioning mechanism being key to Whi5’s role as one of multiple inputs into G1 size control.

In addition to Whi5, there are several other key regulators of *Start*, including the SBF transcription factor, that could contribute to the size-dependence of G1 (Andrews and Herskowitz, 1989; Eser et al., 2011; Ferrezuelo et al., 2010; Koch et al., 1993; Nasmyth and Dirick, 1991). Crucially, SBF regulates the transcriptional activation of G1 cyclins that complete the positive feedback loop defining *Start* as the commitment point to enter the cell cycle (Doncic et al., 2011; Skotheim et al., 2008). SBF was previously identified as a key regulator of super-scaling gene expression in a study examining gene expression in cells of different size in G1 phase (Chen et al., 2020). However, in this data, Whi5 was diluted in the larger G1 cells so that it was possible to interpret the SBF super-scaling to be due to the size-dependent Whi5 dynamics. Interestingly, we found here that the transcription of a number of SBF targets, including the G1 cyclin *CLN2*, super-scales throughout the cell cycle. This is unlikely to be a downstream effect of Whi5 sub-scaling because by this stage in the cell cycle Whi5 has been phosphorylated, inactivated, and exported from the nucleus. Thus, it appears likely that Whi5 is not the only size-dependent signal regulating SBF activated transcription. We find it intriguing that both the super-scaling and sub-scaling genes we identified here are predominantly cell cycle regulated transcripts whose expression peaks outside of G1.

### The role of Whi5 dilution in G1

It is important to note that our Whi5 dilution model does not propose to explain exactly why any given cell enters the cell cycle precisely when it does. Instead it addresses the question of how cells measure their size and then input this information, alongside other size-independent inputs, into the decision to divide. Importantly, the relationship between size at birth and G1 duration exhibits significant cell-to-cell variability (Di Talia et al., 2007). Our model is therefore not that cells progress through *Start* at a precise Whi5 concentration threshold. Rather, our model is that the stochastic rate of progression through *Start* is modulated by the size-dependent Whi5 concentration, as well as additional cell size-dependent and cell size-independent mechanisms. Moreover, we support the recent conclusion that the relative importance of Whi5 sub-scaling and dilution will vary significantly in different growth conditions (Qu et al., 2019). For instance, G1 is longer and cell size control is more pronounced in daughter cells born in poorer carbon sources (*e.g.*, non-fermentable carbons such as ethanol and glycerol) compared to those born in richer carbon sources (*e.g.*, glucose). It is also under poorer carbon sources that Whi5 concentrations are highest (Qu et al., 2019). Thus, daughter cells born in rich carbon conditions will have lower Whi5 concentrations combined with a shorter G1 which results in less Whi5 dilution simply because the extent of dilution is determined by the amount of volume growth in G1. Consistent with this picture, it is precisely in these rich conditions that size control, measured as the degree of inverse correlation between G1 duration and cell size at birth, is weakest (Di Talia et al., 2007).

### Chromatin binding provides an elegant mechanism for equal protein segregation independent of daughter cell size

When examining the inheritance of Whi5 protein levels during cell division we discovered that the inherent asymmetry at cytokinesis poses a problem for sub-scaling proteins. If a protein were simply partitioned in proportion to the relative volumes of newborn cells, sub-scaling would be lost. Instead, we found cells use chromatin-binding to harness the faithful segregation of sister chromosomes to partition approximately equal protein amounts to newborn cells regardless of their size (Fig. 7). This partitioning mechanism is especially relevant when there is a major size asymmetry at cytokinesis as is the case for budding yeast, but also in some metazoan cell divisions including *D. melanogaster* neuroblasts and *C. elegans* cells in the early embryo (Chia et al., 2008; Sulston et al., 1983). Thus, while it has long been appreciated that big and small cells both have the same amount of DNA, here we identified a set of genes that are similarly sub-scaling. It is both curious and elegant that the sub-scaling gene set includes *WHI5*, which regulates DNA replication, while the DNA itself is used as a scaffold for the synthesis and maintenance of sub-scaling protein concentrations.

**Figure S1.**
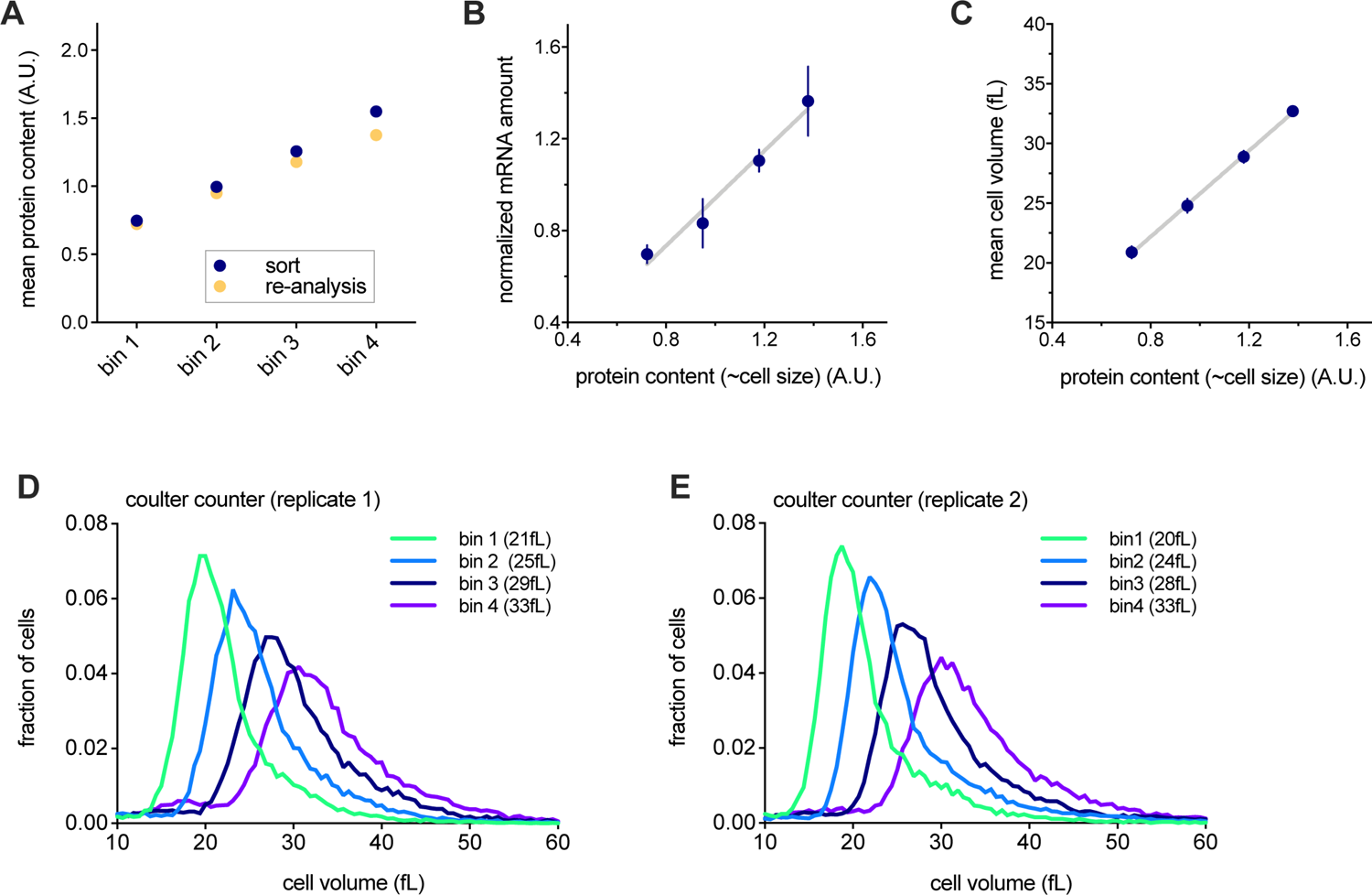
Transcriptomic analysis of cells sorted by cell size. Related to Figures 1D-F. (A-E) Cells in S/G2/M were sorted into four different bins based on the intensity of a total protein dye that we used as a proxy for cell size. See Materials and Methods for further details. (A) Protein dye intensity (normalized to the mean) measured during sorting (blue) and re-measured for cells from each bin after the sort (yellow). The re-analysis (yellow) confirms high sort fidelity. (B) The amount of mRNA (mean ± SEM) per cell for each bin, determined by the number of reads relative to those from a fixed number of *S. pombe* spike-in cells added to the sample. Total protein amount is well correlated with total cellular mRNA amount. (C-E) Cell volume of cells after sorting, measured by Coulter counter. Total protein amount is correlated with cell volume.

**Figure S2.**
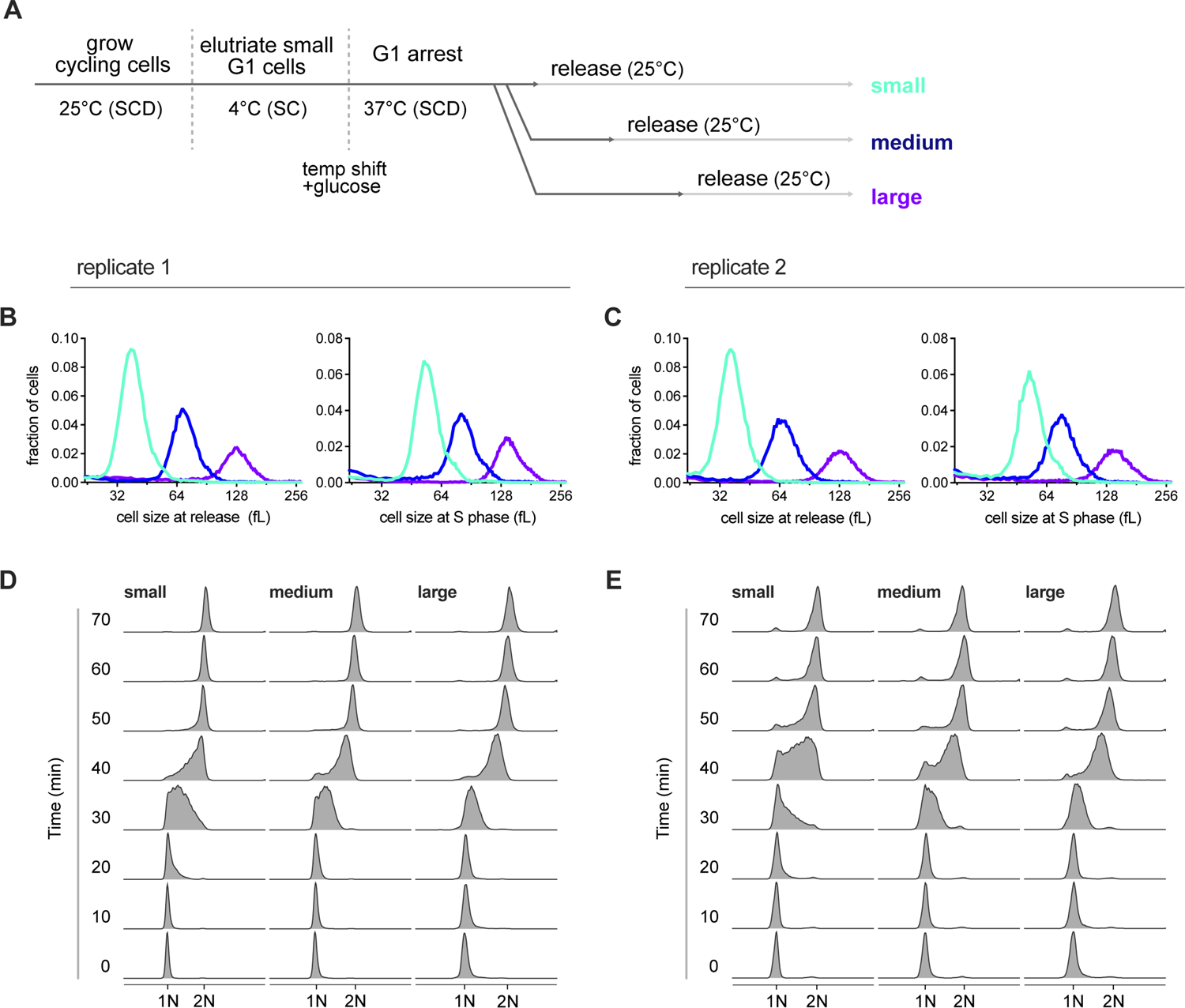
Transcriptomic analysis of different sized cells synchronously progressing through the cell cycle. Related to Figure 2. (A) Schematic of experimental design. G1 cells harboring a temperature sensitive Cdk1 allele (*cdc28-13*) were recovered by centrifugal elutriation. Cells were then shifted to the restrictive temperature (37°C) to arrest them in G1. After increasing amounts of time during the arrest, cells reached small, medium or large cell sizes. The culture was then shifted to the permissive temperature (25°C) to allow cell cycle entry. Samples for RNA-seq were then collected every ten minutes at the eight time points corresponding to synchronous progression through the cell cycle. The 0-minute time point is designated as that 30 minutes before early S phase as determined by DNA-content analysis using flow cytometry (Fig. S2D&E). See Materials and Methods for further details. (B&C) Cell size distributions, measured by Coulter counter, for small, medium, and large cells when they are released from the G1 arrest (left panel) or at mid S-phase (the 40-minute time point, right panel) for two biological replicates. (D&E) DNA content analysis of small, medium, and large cells synchronously progressing through the cell cycle for two biological replicates. Data in Figures S2B&D are also presented in Figure 2A&B (Shown here for comparison with S2C&E).

**Figure S3.**
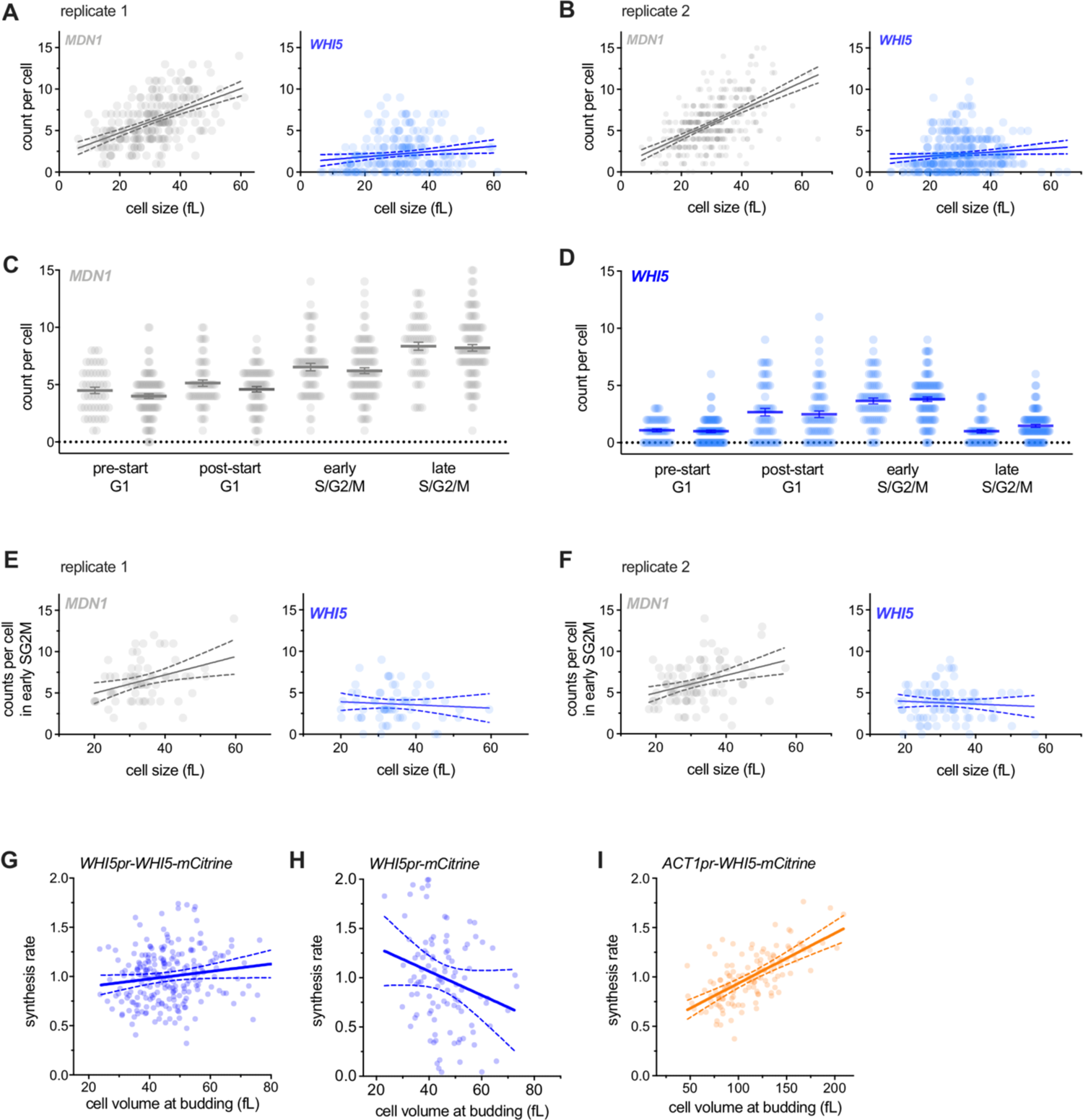
Single-cell analysis of *WHI5* mRNA expression. Related to Figures 1G-I & 2E. (A&B) mRNA counts per cell, measured by single molecule FISH, for *MDN1* or *WHI5* as a function of cell size in two independent biological replicates. All cells of any cell cycle stage are included. Linear regression (solid line) and 95% confidence interval (dashed lines) are shown. The same data are re-plotted in Fig. 1 H&I with replicates pooled. (C&D) mRNA counts per cell, measured by single molecule FISH, for *MDN1* or *WHI5* at different cell cycle stages in two independent biological replicates. Mean (solid line) ± SEM (error bars) are shown. See Materials and methods for details of cell cycle stage classification. (E&F) mRNA counts per cell, measured by single molecule FISH, for *MDN1* and *WHI5* as a function of cell size during early S/G2/M in two independent biological replicates. Early S/G2/M cells were defined as budded cells with a small (≤ 0.2) bud-to-mother volume ratio. Linear regression (solid line) and 95% confidence interval (dashed lines) are shown. The same data are re-plotted in Fig. 2E with replicates pooled. (G-I) Protein synthesis rates (normalized to the mean) as a function of cell volume at budding, measured by time-lapse microscopy for Whi5-mCitrine fusion proteins expressed from (G) the endogenous *WHI5* promoter or (I) the *ACT1* promoter and (H) mCitrine expressed alone from the *WHI5* promoter. Synthesis rates were determined for single cells for the period between bud emergence and cytokinesis using a linear fit as in (Schmoller et al., 2015). Linear regression (solid line) and 95% confidence interval (dashed lines) are shown. (G) n=229, 2 data points outside the axis limits. (I) n=113, 4 data points outside the axis limits. (H) n=132. The same data, binned by cell volume at budding, are shown in Fig. 2F-H.

**Figure S4.**
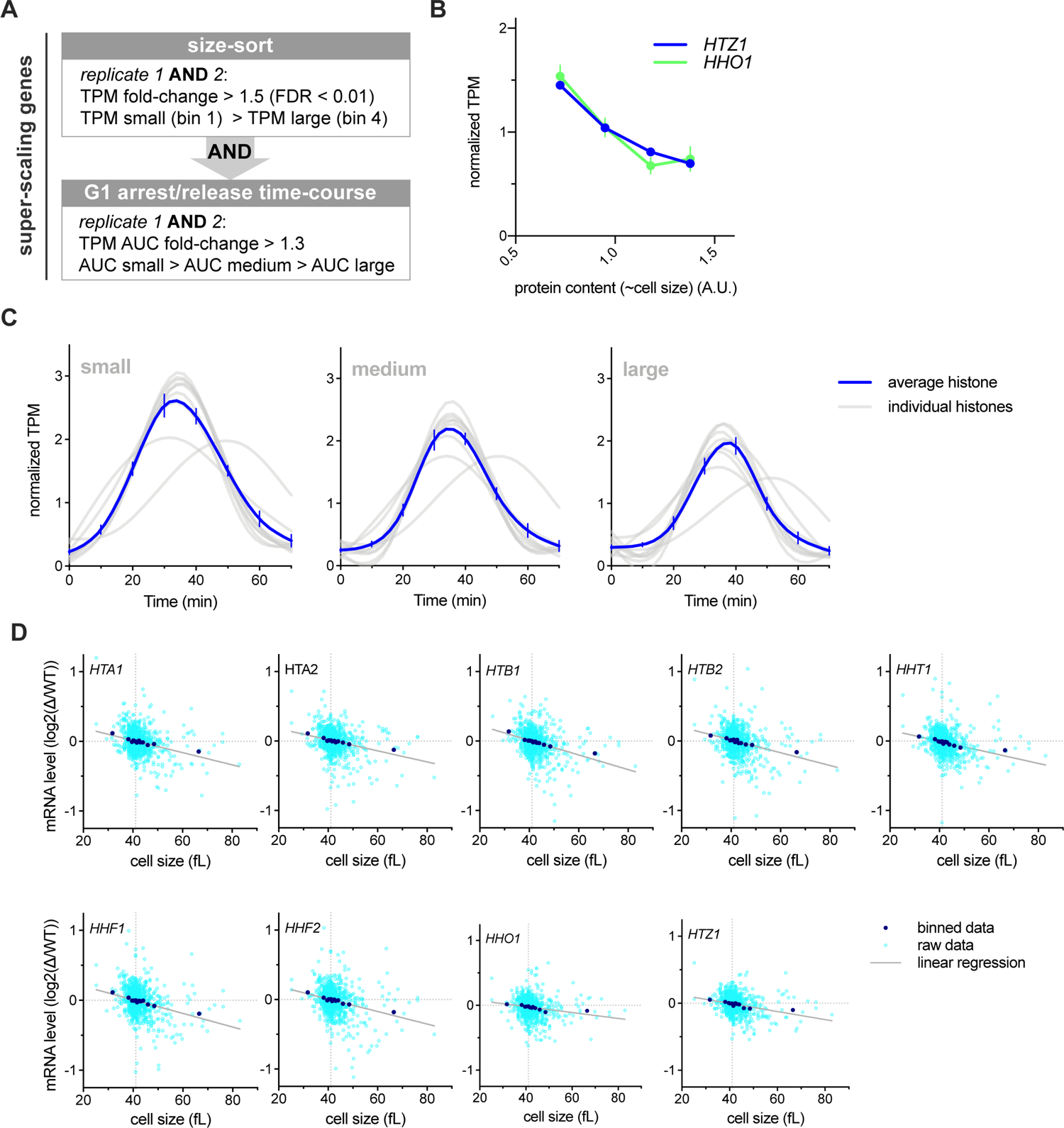
Histone gene expression sub-scales with cell size. Related to Figure 3. (A) Criteria used to classify genes with sub-scaling transcript amounts. Genes whose expression changed in a manner similar to *WHI5* in both replicates of the size-sort (see Fig. 1D-F) and in the G1 arrest/release time course (see Fig. 2A-D) were classified as sub-scaling genes. See Materials and Methods for further details. (B) Normalized TPM (TPM / mean TPM) for *HTZ1* and *HHO1* mRNA in cells of different sizes (total protein content). Mean (± range) of two biological replicates is plotted. Changes in TPM are proportional to changes in mRNA concentration. See Figure S1 and Materials and Methods for further details. (C) Normalized histone mRNA TPM (TPM / mean TPM) of small, medium or large cells at 10-minute time intervals synchronously progressing through the cell cycle. A smoothing spline fitted to the mean normalized TPM from two biological replicates for each histone is plotted in grey. A Spline fitted to mean (± SEM) of all histones is shown in blue. (D) Comparison of histone mRNA levels in 1,484 gene deletions, relative to wild-type (Kemmeren et al., 2014; O’Duibhir et al., 2014) with cell size of the respective gene deletions (Jorgensen et al., 2002). Each graph plots a single transcript and each point represents a single gene deletion. See Materials and Methods for details. Distribution of Pearson R correlation coefficients for all genes in is show in Fig. 3F.

**Figure S5.**
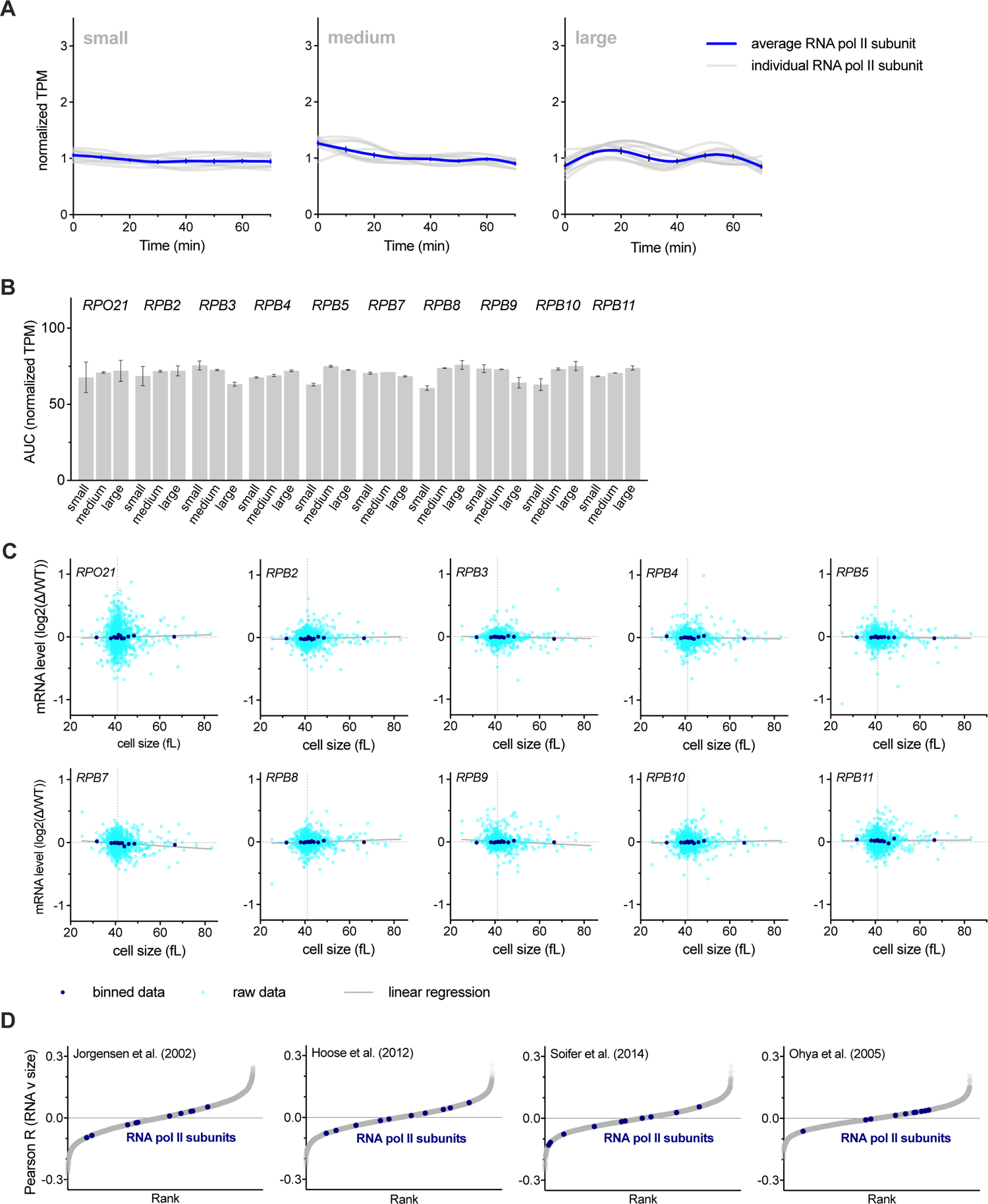
RNA polymerase II subunit gene expression scales and is proportional to cell size. Related to Figure 3 (A) Normalized RNA polymerase II subunit mRNA TPM (TPM / mean TPM) for small, medium, and large cells at 10-minute time intervals synchronously progressing through the cell cycle. A smoothing-spline fitted to the average normalized TPM from two biological replicates for each subunit is plotted in grey. A Spline fitted to the mean (± SEM) of all subunits is shown in blue. (A) (B) The Area Under the Curve (AUC) of mean normalized RNA polymerase II subunit mRNA TPM of small, medium, and large cells synchronously progressing through the cell cycle. The AUC mean (± range) of two biological replicates is plotted. (B) Comparison of RNA polymerase II subunit mRNA levels in 1,484 gene deletions, relative to wild-type (Kemmeren et al., 2014; O’Duibhir et al., 2014) with cell size of the respective gene deletions (Jorgensen et al., 2002). Each graph plots a single transcript and each point represents a single gene deletion. See Materials and Methods for details. (C) Pearson correlation coefficient, R, for the correlation between mRNA levels relative to wild-type in 1,484 gene deletion strains (Kemmeren et al., 2014; O’Duibhir et al., 2014) and the cell size of the respective gene deletions for four different data sets of size measurements (Hoose et al., 2012; Jorgensen et al., 2002; Ohya et al., 2005; Soifer and Barkai, 2014). Each point represents an individual gene. RNA polymerase II subunit mRNAs are shown in blue.

**Figure S6.**
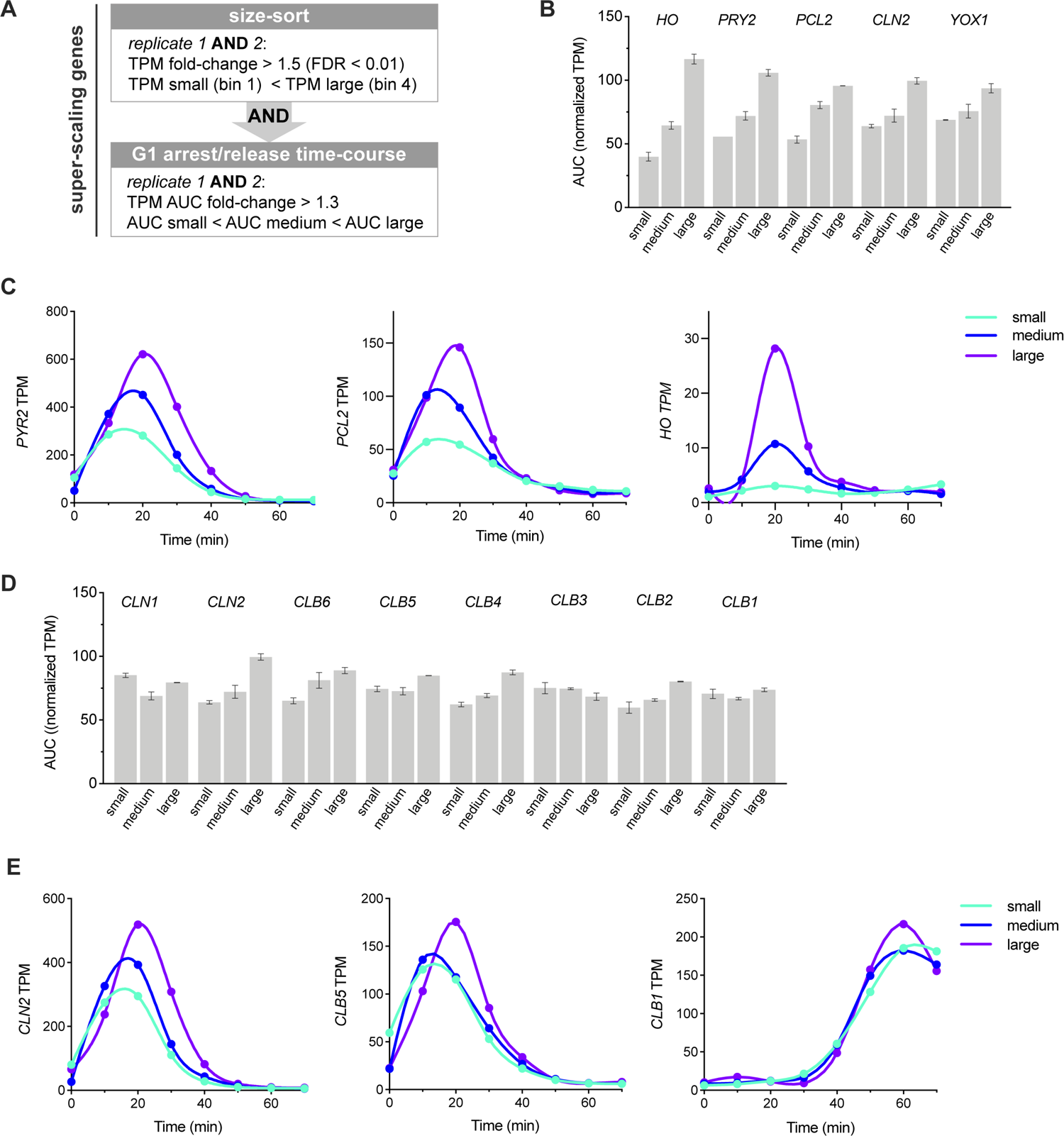
SBF regulated super-scaling genes and cyclin expression during the cell cycle in cells of different sizes. Related to Figure 3. (A) Criteria used to classify genes with super-scaling transcript amounts. Genes whose expression increased in both replicates of the size-sort and in the G1 arrest/release time course were classified as super-scaling genes. See Materials and Methods for further details. (B) The Area Under the Curve (AUC) of mean normalized TPM of small, medium, and large cells synchronously progressing through the cell cycle for SBF-regulated super-scaling transcripts. See Fig. S2 for experimental details. The AUC mean (± range) of two biological replicates is plotted. (C) SBF-regulated super-scaling genes TPM for small, medium, and large cells synchronously progressing through the cell cycle. Data for one replicate is plotted and is representative of both replicates. (D) The Area Under the Curve (AUC) of mean normalized TPM of small, medium, and large cells synchronously progressing through the cell cycle for cyclin transcripts. The AUC mean (± range) of two biological replicates is plotted. (E) Cyclin genes TPM for small, medium, and large cells synchronously progressing through the cell cycle. See Fig. S2 for experimental details. Data for one replicate is plotted and is representative of both replicates.

**Figure S7.**
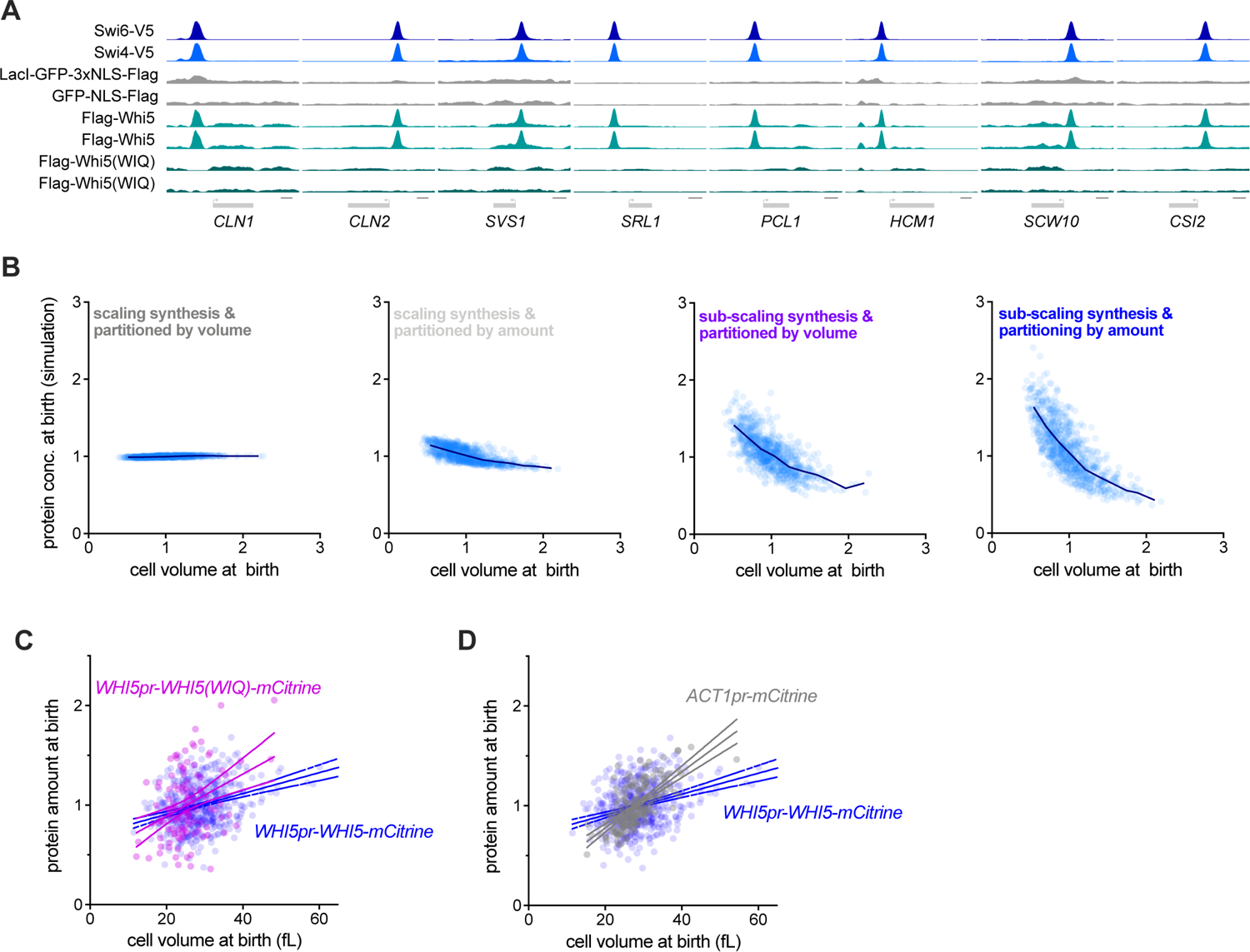
Sub-scaling expression is inherited through asymmetric division due to DNA-mediated partitioning. Related to Figure 5. (A) anti-V5 or anti-Flag ChIP-seq experiments were performed using cells expressing the indicated fusion proteins (left hand side). Data are shown for 8 example SBF binding sites near the denoted genes. Cells expressing LacI-GFP-3xNLS-FLAG or GFP-NLS-FLAG were included as controls for non-specific ChIP signal. Whi5 recruitment to DNA overlaps with SBF (Swi4 and Swi6) binding sites but is lost in the Whi5(WIQ) variant (Travesa et al., 2013). Summary metagene plots for a subset of these data are also shown in Fig. 5C. (B) Simulation protein concentration at birth as a function of daughter volume at birth. Four different conditions were simulated where protein expression was either in proportion to cell size (scaling) or independent of cell size (sub-scaling), and protein partitioning is either by amount or in proportion to cell volume. Individual simulated cells (light blue) as well as bin means (dark blue) are plotted. Protein amounts from the same simulation are shown in Fig. 5D. (C&D) Protein amount at birth for *WHI5pr-WHI5-mCitrine* and (C) *WHI5pr-WHI5(WIQ)-mCitrine* or (D) *ACT1pr-mCitrine* cells. Linear regression (solid line) and 95% confidence interval (dashed lines) are shown. Note, Fig. 5E shows bin means of the same data as (C) binned by cell volume at birth.

**Figure S8.**
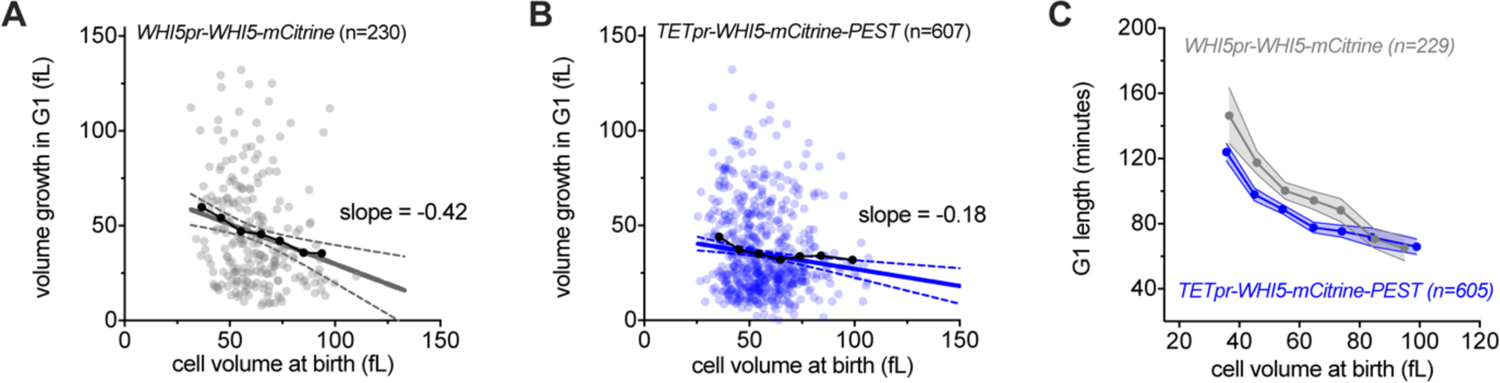
Expression of Whi5-PEST fusion protein from a synthetic TET-inducible promoter attenuates G1 size control. Related to Figure 6 (A&B) Cell volume growth in G1 (birth to budding) plotted against cell volume at birth for (A) *WHI5pr-WHI5-mCitrine bck2Δ* and (B) *TETpr-WHI5-mCitrine-PEST bck2Δ.* Linear regression (solid line) and 95% confidence interval (dashed lines) are shown. Linear regressions are calculated after outlier removal (Q=1%), which resulted in 2 (A) and 7 (B) cells being excluded respectively. Note, Fig. 6D shows bin means of the same data binned by cell volume at birth. (C) Time in G1 as a function of cell size at birth, for cells with a completed G1. Data are binned according to cell size at birth for cells with a completed G1 and the bin means (± SEM) are plotted.

**Table S1.** List of *S. cerevisiae* strains used in this study and full genotypes

|  |  |  |
| --- | --- | --- |
| A17896 | Amon lab | W303; MATa, ADE2, cdc28-13 |
| DCB99 | This study | W303; MATa, ADE2, WHI5-mCitrine::URA3 |
| MK551-1 | This study | W303; MATa, bar1Δ::HisG, whi5Δ::LEU2, cln3Δ::hphMX |
| HTA1-GFP | GFP collection<br>(Huh et al., 2003) | BY4741; HTA1-GFP |
| HTB2-GFP | GFP collection<br>(Huh et al., 2003) | BY4741; HTB2-GFP |
| HTZ1-GFP | GFP collection<br>(Huh et al., 2003) | BY4741; HTZ1-GFP |
| RPB3-GFP | GFP collection<br>(Huh et al., 2003) | BY4741; RPB3-GFP |
| DCB72 | This study | W303; MATa, ADE2, ura3:: WHI5pr(1kb)-mCitrine-CYC1term |
| KSY108-1 | Lab collection | W303; ADE2, WHI5-mCitrine-HIS3 |
| KSY190-2 | This study | W303; ADE2, whi5Δ::KanMX URA3::WHI5pr(1kb)-WHI5-WIQ-mCitrine-ADH1term |
| KSY160-2 | This study | W303; ADE, URA3::ACT1pr(1kb) -Whi5-mCitrine-CYC1term |
| KSY158-1 | Lab collection | W303; ADE2, URA3:: ACT1pr(1kb)-mCitrine-CYC1term |
| MS534 | This study | W303; bar1Δ::HisG, GFP-3xNLS-FLAG |
| MS535 | This study | W303; bar1Δ::HisG, LacI-GFP-3xNLS-FLAG |
| MS536 | This study | W303; bar1Δ::HisG, whi5Δ::LEU2, URA3::WHI5pr(1kb)-3xFLAG-WHI5-CYC1term |
| MS537 | This study | W303; bar1Δ::HisG, URA3::WHI5pr(1kb)-3xFLAG-WHI5(WIQ)-CYC1term |
| MK653-1 | This study | W303; bar1Δ::HisG, SWI4-V5::hphMX6 |
| MK645-1 | This study | W303; bar1Δ::HisG, SWI6-V5::hphMX6 |
| M364 | This study | W303; whi5Δ::cgTRP; bck2Δ::hphNT, myo1-3xmKate2::kanMX6, leu1::PtetO7.1-WHI5-mCitrine-PEST::LEU1, WTC846(PrtetO-7.1-TetR, PrRNR2-TetR-TUP1)::HIS3, ADE |
| M367 | This study | W303; whi5Δ::cgTRP; bck2Δ::hphNT1, myo1-3xmKate2::kanMX6, WHI5pr[1kb]-WHI5-mCitrine::URA3, WTC846(PrtetO-7.1-TetR_PrRNR2-TetR-TUP1)::HIS3, ADE |

## Acknowledgments

We thank Bruce Futcher and Yuping Chen for invaluable advice on centrifugal elutriation, Christine Jacobs-Wagner and Helena Cantwell for comments on the manuscript, members of the Skotheim laboratory for constructive feedback, Gabriel Neurohr for sending strain A17896, Chris You for assistance with the ChIP-seq experiments and Jon Turner for help in optimizing the smFISH protocol. This work was supported by the NIH (GM092925 and GM115479), the HHMI-Simons (JMS, Faculty Scholars Program). MPS was supported by a Simons Foundation Fellowship of the Life Sciences Research Foundation and an EMBO Long-Term Postdoctoral Fellowship. KMS was supported by the Human Frontier Science Program (Postdoctoral Fellowship and Career Development Award).

## Author contributions

JMS supervised the work. MPS and JMS wrote the manuscript. MPS performed and analyzed all experiments except for live cell microscopy experiments performed by JK, DCB & KMS, and smFISH experiments performed by ML. Sequencing data analysis was performed by MPS and GM, supervised by WG & AK.

## MATERIALS AND METHODS

### Yeast genetics

Standard procedures were used for *Saccharomyces cerevisiae* strain construction. Full genotypes of all strains used in this study are listed in Table S1. The strain for constitutive expression of WHI5 (MS364) was constructed using the WTC846 tetracycline responsive promoter system (Azizoğlu et al., 2020).

### Transcriptomic analysis of size-sorted cells

To determine transcript levels in cells of different sizes, S/G2/M cells were sorted according to total protein content by fluorescence activated cell sorting (FACS) before RNA extraction and sequencing. 500ml *S. cerevisiae* (*HTB2-GFP*) was grown (synthetic complete media + 2% glucose at 30°C), and fixed at O.D. ∼ 0.3 by addition of 500ml 80% methanol 20mM TRIS (−20°C) and then incubated at −20°C for 30 minutes. Cells were fixed to prevent gene expression changes during the course of the cell sorting, which requires multiple generation equivalents of time to complete. Cells were pelleted (13 krpm, 3 minutes) and washed 3x in PBS, before gentle sonication and then addition of 5µg/ml total protein dye (Alexa Fluor™ 647 NHS Ester dye; ThermoFisher Scientific; A20006) and incubation (4°C, 30 minutes). Cells were again pelleted (13 krpm, 3 minutes) and washed 3x in PBS to remove excess dye before again being sonicated. Cells from four different size fractions were sorted on a FACSAria II sorter (BD Biosciences) according to the following strategy. First, singlets were gated based on scatter (FSC and SSC), then S/G2/M cells were identified using an Htb2-GFP signal, and then finally, four bins of different total protein content cells were sorted based on the total protein intensity (bin 1 = lowest signal, bin 4 = highest, Fig. S1A). The fidelity of the total protein dye sort was confirmed by re-analyzing 10,000 cells on the same sorter (Fig. 1D & S1A). Furthermore, total protein content was validated as a proxy for size by measuring the cell volume of the different protein dye sorted cells using a Coulter counter (Fig. S1C-E).

Two biological replicates were performed. Within each biological replicate, two technical replicates were performed for each size bin so that four replicates were performed per bin in total. For the different bins within the same replicate set, a constant number of *S. pombe* cells, fixed as above, were added as a spike-in to measure total RNA content per *S. cerevisiae* cell. The number of *S. cerevisiae* cells and *S. pombe* cells (972 h-) per sample was constant within a set of replicates but varied slightly between each set of replicates (5-10 million *S. cerevisiae* cells and 5-10 million *S. pombe* cells). To maximize the number of reads from the experimental *S. cerevisiae* samples, *S. pombe* cells were nitrogen starved (grown in EMM before media switch to EMM - NH4Cl for 24 hours at 30°C) because this reduces their mRNA concentration (Marguerat et al., 2012). Cells were then pelleted (4 krpm, 15 minutes), and their RNA extracted and sequenced as described below. For each set of replicates, the *S. pombe* spike-in was added independently and RNA was extracted independently.

To estimate the relative amount of mRNA per cell in each size bin, the number of *S. cerevisiae* reads per *S. pombe* read was calculated (see RNA-seq data processing below) and then normalized to the mean value within a set of replicates. The normalized total mRNA per sample was then averaged between the four replicates (Fig. S1B).

### Transcriptomic analysis of differently sized cells synchronously progressing through the cell cycle

To determine transcript levels during the cell cycle in cells of different sizes, cells were elutriated and arrested in G1 for different amounts of time. Samples were then collected during the synchronous progression from G1 through S, G2 and M phases of the cell cycle for RNA extraction and sequencing (See Fig. S2 for schematic of experimental design). Specifically, 4L *S. cerevisiae* (A17896: *W303 cdc28-13*) were grown in synthetic complete (SC) media with 2% glucose at 25°C to OD ∼0.75 and then collected on a filter membrane and resuspend in ice-cold SC media (no carbon source). Cells were then sonicated (3 x 20 seconds, 3 minutes on ice between sonication cycles) and loaded into a JE 5.0 elutriation rotor fitted for a two-chamber run (Beckman Coulter) in a J6-MI Centrifuge (2.4krpm, 4°C). The elutriation chambers were pre-equilibrated and run with SC media (4°C, no carbon source). The pump speed was gradually increased until G1 cells with minimal debris were collected. G1 fractions were then collected on a filter and resuspended in 37°C conditioned SC media + 2% glucose in a 37°C shaking water-bath (OD ∼ 0.1). The G1 arrest was maintained at 37°C until cells reached either 36-39 fL (small), 67-69 fL (medium) or 129-131 fL (large) as determined by Coulter counter (Fig. S2B-C). When they reached the target size, cells were released from the G1 arrest. To do this, cells were collected on a filter membrane and resuspended in 25°C SC media + 2% glucose (OD ∼0.35). Samples for size measurement by Coulter counter, DNA-content analysis, and RNA-extraction were taken at 10-minute intervals after release with the 0-minute time point being designated as the time point 30 minutes before the onset of DNA replication (Fig. S2D-E). For small cells, the 0-minute time point was collected 40-50 minutes after the shift to the permissive temperature, for medium cells the 0-minute time point was collected 10 minutes after the shift to the permissive temperature and for large cells the 0-minute time point was collected 0 minutes after shift to the permissive temperature. Two biological replicates were performed.

For DNA-content analysis, 0.4 mL culture was added to 1 mL 100% 4°C ethanol and stored at 4°C. Cells were pelleted (13 krpm, 2 minutes), washed, and resuspended in 50 mM Sodium Citrate (pH = 7.2), incubated with 0.2 mg/mL RNAse A (overnight, 37°C) and then 0.4 mg/mL proteinase K (1 hour, 50°C) before addition of 25 μM Sytox Green (ThermoFisher Scientific). Cells were then sonicated and DNA-content was analyzed for >10000 events on a FACScan Analyzer (BD Biosciences). For RNA-extraction 1.5 mL cells were pelleted (13 krpm, 30 seconds) and snap frozen in liquid N_2_. Samples were then thawed in TRI Reagent (Zymo Research) and RNA was extracted as described below (RNA extraction and sequencing).

### RNA extraction and sequencing

To extract RNA, cell pellets were lysed in 300 µL TRI Reagent (Zymo Research) by bead beating using a Fastprep 24 (4°C, settings: 5.0 m/s, 1 x 30 seconds). Cell debris was pelleted (13 krpm, 5 minutes) and the supernatant recovered. RNA was then extracted using the direct-zol RNA microprep kit (Zymo Research). mRNA was enriched using the NEBNext Poly(A) mRNA Magnetic Isolation Module (NEB, E7490) and NEBNext Ultra II RNA Library Prep Kit for Illumina^®^ (NEB, #E7775) was then used to prepare libraries for paired-end (2×150 bp) Illumina sequencing (Novogene). More than 20 million reads were sequenced per sample.

### RNA-seq data processing

Because some samples analyzed in this study contained *S. cerevisiae* as well as reference spike-in *S. pombe* RNA, a combined *S. cerevisiae* and *S. pombe* genome file was created using the sacCer3 and ASM294v2 versions of the respective genomes and a combined transcriptome annotation was created using the *S. pombe* gene models available from PomBase (Lock et al., 2019) and an *S. cerevisiae* set of gene models updated using transcript-end mapping data as previously described (Shipony et al., 2020). For the purposes of RNA-seq data quality evaluation and genome browser track generation, reads were aligned against the combined genome and annotated set of splice junctions using the STAR aligner (version 2.5.3a; settings: --limitSjdbInsertNsj 10000000 -- outFilterMultimapNmax 50 --outFilterMismatchNmax 999 --outFilterMismatchNoverReadLmax 0.04 -- alignIntronMin 10 --alignIntronMax 1000000 --alignMatesGapMax 1000000 --alignSJoverhangMin 8 --alignSJDBoverhangMin 1 --sjdbScore 1 --twopassMode Basic --twopass1readsN −1) (Dobin et al., 2013). Read mapping statistics and genome browser tracks were generated using custom Python scripts. For quantification purposes, reads were aligned as 2×50mers in transcriptome space against an index generated from the combined annotation described above using Bowtie (Langmead et al. 2009; version 1.0.1; settings: -e 200 -a -X 1000). Alignments were then quantified using eXpress (version 1.5.1) (Roberts and Pachter, 2013) before effective read count values and TPM (Transcripts Per Million transcripts) were then separated for each genome and renormalized TPMs were calculated with respect to the total reads for *S. cerevisiae*.

Differential expression analysis by DESeq2 was performed using technical replicates to compare RNA-seq data from different size bins in the experiment shown in Figures 1D-F and S1 (Love et al., 2014). To calculate the total amount of transcription during the G1 arrest/release RNA-seq time course experiment (Fig. 2C-D & S2), TPM values were normalized to the mean for each experiment and the Area Under the Curve (AUC) for TPM as a function of time was calculated for each time course using the R function auc(*type = “spline”*) from the *MESS* package.

### Classification of sub-scaling and super-scaling transcripts

To classify transcripts whose expression sub-scales with cell size, we analyzed data from two experiments: (1) the RNA-seq experiment on size-sorted populations of cells (Fig. 1D-F and S1) and (2) the G1 arrest/release RNA-seq time course experiment (Fig. 2A-D and S2). Two biological replicates of each experiment were performed. Sub-scaling genes were classified as genes that passed the following criteria in both biological replicates of each experiment:

1. At least one pair-wise comparison between the four-size bins has a false-discovery rate (FDR) adjusted p-value <0.01, a TPM fold-change > 1.5, and bin 1 TPM > bin 4 TPM.
2. Small cells’ TPM Area Under Curve (AUC) > medium cells’ TPM AUC > large cells’ TPM AUC and TPM AUC fold-change > 1.3. See above (RNA-seq data processing) for details of the AUC calculation.

To classify transcripts whose expression sub-scales with cell size we applied same criteria as for sub-scaling genes but instead (1) bin 1 PM < bin 4 TPM and (2) Small cells’ TPM Area Under Curve (AUC) < medium cells’ TPM AUC < large cells’ TPM AUC.

### GO term enrichment

GO term enrichment (Fig. 3A) was performed using the GOrilla GO analysis tool (Eden et al., 2009). Enrichment of size-independent gene transcripts was performed versus a background set of all genes that had a TPM value > 0 in all RNA-seq samples.

### Single-molecule fluorescence in situ hybridization (smFISH)

smFISH was used to image *WHI5* and *MDN1* mRNAs in single cells. A Whi5-mCitrine tagged strain (DCB099) was used to discriminate pre- and post-*Start* G1 based on Whi5-mCitrine nuclear localization (Doncic et al., 2011). Early S/G2/M cells were defined as budded cells with a small (≤ 0.2) bud-to-mother volume ratio. Cells were grown at 30°C in synthetic complete (SC) media + 2% glycerol + 1% ethanol. Two biological replicates were performed. Each biological replicate contained two technical replicates (*i.e.*, two independent hybridizations to cells from the same culture). In addition, two negative controls were performed regularly where (i) FISH probes were omitted and (ii) a *whi5Δ* (MK551-1) strain was analyzed.

The smFISH protocol (detailed below) was optimized based on protocols from multiple prior studies (Raj and Tyagi, 2010; Trcek et al., 2012; Tutucci et al., 2018; Youk et al., 2010; Zenklusen et al., 2008). 45 mL cells (OD600 ∼0.2) were fixed with 5 mL 37% formaldehyde and incubated (45 minutes, room temperature, rotating). Cells were then pelleted (1600 g, 5 minutes) and washed twice in 1 mL of ice-cold fixation buffer (pH 7.5, 218 mg/mL sorbitol, 84 mM potassium phosphate dibasic, 16 mM potassium phosphate monobasic, dissolved in water for RNA work (Thermo Fisher Scientific, BP561-1)). Cells were again pelleted and resuspended in 900 μL fixation buffer and gently sonicated before 100 μL of 200 mM RNase inhibitor vanadyl ribonucleoside complex was added (New England BioLabs, S1402S). Cells were then digested by adding 3.5-5 μL zymolyase stock (5 mg/mL 100T, MP Biomedicals, 0832093) and incubated (70-80 minutes, 30°C, rotating). Cells were then pelleted (400 g, 6 minutes) and washed twice in 1 mL of ice-cold fixation buffer to stop digestion and finally permeabilized by resuspension in 1ml 70% ethanol. Permeabilized cells were kept at 4°C for 1 to 3 days. 300 μL of the permeabilized cells per hybridization sample were then pelleted (400 g, 7 minutes) and washed in 500 μL of wash buffer A (Biosearch Technologies, SMF-WA1-60: prepared fresh on the day of use according to manufacturer’s instructions always using a fresh aliquot of deionised formamide (EMD Millipore, S4117, stored at −20°C)). Permeabilized cells were then resuspended in 100 μL hybridisation solution (Biosearch Technologies, SMF-HB1-10) containing 1-3x of standard probe concentrations for *WHI5* and *MDN1* probes, 10 mM VRC and 0.5 mg/mL smFISH probe competitor E. coli tRNA (Roche, TRNAMRE-RO). Note VRC and probe competitor were omitted for half the cells analyzed in replicate 1. For *MDN1* mRNAs two probe sets, totaling 86 probes (38 + 48), coupled to the FAM (fluorescein amidite) dye (Biosearch Technologies, Stellaris Custom Probes) were used. The sequences of these probes were taken from Tutucci et al., 2018 (*MDN1*-3’ORF and *MDN1*-ORF). For *WHI5*, a set of 46 probes coupled to the Quasar570 dye (Biosearch Technologies, Stellaris Custom Probes) were used. *WHI5-mCitrine* Probe sequences were designed using Stellaris Probe Designer applied to the *WHI5-mCitrine* mRNA sequence. Probes were hybridized in the dark (30°C, overnight, with end-over-end rotation). 100 μL of wash buffer A was then added before cells were pelleted (400 g, 8 min) and supernatant was aspirated. Cells were resuspended in 1 mL wash buffer A, incubated in the dark (30° C, 30 min), pelleted (400 g, 6 min), resuspended in 1 mL wash buffer A +350 μg/mL calcofluor white (Sigma, F3543), and again incubated in the dark (30°C, 30 min). Cells were then resuspended in 1 mL wash buffer B (Biosearch Technologies, SMF-WB1-20), incubated for 2-5 minutes at room temperature, pelleted (400 g, 6 minutes) and resuspended in 2 to 3 drops (∼75 μL) of Vectashield Antifade Mounting Medium (Vector Laboratories, H-1000). This suspension was then mixed thoroughly by pipetting to separate clumped cells. 1.5 μL of this solution was mounted on an acid washed slide and imaged on a wide-field epifluorescence Zeiss Observer Z1 microscope (63X/1.4NA oil immersion objective and a Colibri LED module). 30-step *z*-stacks (step size = 200 nm) were imaged. Cell outlines were identified using phase contrast images. Quasar570 probes (*WHI5*) were imaged in the orange channel (white LED module for 555 nm wavelength, 100% light power, 5 sec exposure per stack image). *Whi5-mCitrine* protein and FAM probes (*MDN1*) were imaged in the yellow channel (505 nm LED module, 100% light power, 3.5 seconds exposure time). FAM FISH probes alone were imaged in the green channel (470 nm LED module, 75% light power, 5 seconds exposure time). Calcofluor white stain was imaged in the blue channel (365 nm LED module, 25% light power, 20 ms exposure). Under these conditions, no significant photobleaching was observed after taking multiple images of the same cells.

smFISH Image analysis was performed manually using ImageJ (version 2.0.0). Single cells were manually selected in each image. A cell was only selected if its morphology was sufficiently intact (following zymolyase treatment) and if the absence/presence of a bud and the nuclear/cytoplasmic localization of Whi5-mCitrine protein could be assigned. Cell size was measured by drawing cell outlines in the phase *z*-plane with the largest cell area, fitting a two-dimensional ellipse, and then rotating the ellipse along its major axis to obtain a volume estimate. Separately calculated volumes for mothers and buds were added together. Absolute counts of *WHI5-mCitrine* and *MDN1* mRNAs in single cells were obtained by manual counting and single dots were counted as one mRNA (*i.e.*, we did not quantify single dot intensities to try to discern multiple overlapping mRNAs).

### Live cell microscopy

Cells were grown to early log phase in synthetic complete (SC) media + 2% glycerol + 1% ethanol and gently sonicated before being loaded into a CellASIC Y04C microfluidics plate (Milipore SIGMA) under continuous media flow at 2 psi. Imaging, image segmentation, and pedigree tracking was performed as previously described (Doncic et al., 2013; Schmoller et al., 2015) with the exception of the experiments in Fig. 6 which were segmented and tracked using the convolutional neural network YeaZ algorithm (www.quantsysbio.com). For experiments in Fig. 6, Myo1-3xmKate2 signal at the bud neck was also used aid the determination of mother-bud pairs and cytokinesis timing. Cells expressing GFP proteins were exposed for 50 ms, cells expressing mCitrine were exposed for 400 ms and cells expressing mKate2 were exposed for 1s. Background subtraction for variation in background fluorescence in each frame of the movie was performed as previously describe (Chandler-Brown et al., 2017). Briefly, in each frame cell and non-cell area was defined. A 4-pixel average filter was then applied and the background was taken to be the median filtered pixel value of the non-cell area except for experiments in Fig. 6 where background fluorescence was measured using median pixel values from a collection of flatfield images. Differences in background fluorescence due to cell volume dependent autofluorescence were accounted for as previously described (Schmoller et al., 2015). Note that the analysis of *WHI5pr-WHI5-mCitrine* (Fig. 2F, S3G & 5B&E) includes cells previously imaged for analysis reported in Schmoller et al., 2015.

### Gene deletion collection screen by microarray

We analyzed the correlation between RNA levels and cell size in 1,484 gene deletion stains using published microarray data and cell size measurements of these strains (Hoose et al., 2012; Jorgensen et al., 2002; Kemmeren et al., 2014; O’Duibhir et al., 2014; Ohya et al., 2005; Soifer and Barkai, 2014). Gene expression changes relative to wild-type were from (O’Duibhir et al., 2014), where we used the dataset transformed to correct for effects of slow growth. The same trends were observed in the uncorrected dataset. For each gene we then analyzed the correlation between the relative fold-change of its expression in a given deletion with the size of that deletion strain across all the deletion strains for which both data were available. Pearson correlation coefficients were calculated using the R function *cor* (Fig. 3F & S5D).

### GFP fusion collection screen by flow cytometry

To examine the size-dependence of histone protein’s expression, we analyzed a genome-wide dataset of flow cytometry-based GFP intensity measurements (Parts et al., 2014), where each measurement is from a single well containing two strains both expressing the same protein C-terminally fused to GFP (Fig. 4A-C). One strain is in the BY4741 background (replicate 1) and the other in the RM11 background (replicate 2). Cells were grown in low fluorescence media containing 2% glucose and measured using an BD LSRII flow cytometer as described in Parts et al., 2014. Cells were separated into budded and unbudded populations on the basis of the side scatter width (SSC-W). Co-cultured strains of each background were separated on the basis of HTB2-mCherry intensity (RM low, BY high). Size was defined by the area of the side scatter signal (SSC-A). We used the lowest expressed gene from each plate as a background for that plate, thereby controlling for plate-to-plate variation in measurements. To calculate the background, we fitted a linear function to SSC-A and total GFP fluorescence for these low-expressing cells (python function *polyfit*(matplotlib)). We then subtracted the fit for these lowest expressing cells from the GFP intensity for all other cells. Strains with noisy signals (*i.e.*, their mean expression is less than the standard deviation) and cells with saturated signals (mean expression is greater than 200000) were excluded. SSC-A and background subtracted GFP intensity were then normalized to the mean and a linear function was then fitted (python function *polyfit*(matplotlib)). The slope of this function was used as a measurement for a protein’s size-dependence.

### ChIP-seq experiments

Cells expressing Swi4-V5, Swi6-V5, 3xFLAG-WHI5, 3xFLAG-WHI5, GFP-NLS-5xFLAG or LacI-GFP-NLS-5xFLAG were grown in SC media with 2% glycerol 1% ethanol. 500 mL of cells at OD ∼0.5 were fixed with 1% formaldehyde (30 minutes) and quenched with 0.125 M glycine (5 minutes). Fixed cells were washed twice in cold PBS, pelleted, snap-frozen and stored at −80°C. Cell lysis and ChIP reactions were performed as previously described (Hu et al., 2015) with minor modifications. Pellets were lysed in 300 µL FA lysis buffer (50 mM HEPES–KOH pH 8.0, 150 mM NaCl, 1 mM EDTA, 1% Triton X-100, 0.1% sodium deoxycholate, 1 mM PMSF, Roche protease inhibitor) with ∼1 mL ceramic beads using a Fastprep-24 (MP Biomedicals). The entire lysate was then collected and adjusted to 1 mL before sonication with a 1/8’ microtip on a Q500 sonicator (Qsonica) for 15 minutes (10 seconds on, 20 seconds off). The sample tube was held suspended in a −20°C 80% ethanol bath to prevent sample heating during sonication. Cell debris was then pelleted and the supernatant retained for ChIP. For each ChIP reaction, 30 µL Protein G Dynabeads (Invitrogen) were blocked (PBS + 0.5% BSA), prebound with 10 µL anti-V5 antibody (SV5-Pk1, BioRad Cat# MCA1360G) or 10 µL anti-FLAG antibody (M2, SIGMA Cat# F1804) and washed once with PBS before incubation with supernatant (4°C, overnight). Dynabeads were then washed (5 minutes per wash) twice in FA lysis buffer, twice in high-salt FA lysis buffer (50 mM HepesKOH pH 8.0, 500 mM NaCl, 1 mM EDTA, 1% Triton X-100, 0.1% sodium deoxycholate, 1 mM PMSF), twice in ChIP wash buffer (10 mM TrisHCl pH 7.5, 0.25 M LiCl, 0.5% NP-40, 0.5% sodium deoxycholate, 1 mM EDTA, 1 mM PMSF) and once in TE wash buffer (10 mM TrisHCl pH 7.5, 1 mM EDTA, 50 mM NaCl). DNA was eluted in ChIP elution buffer (50 mM TrisHCl pH 7.5, 10 mM EDTA, 1% SDS) at 65°C for 15-20 minutes. Eluted DNA was incubated to reverse crosslinks (65°C, 5 hours), before treatment with RNAse A (37°C, 1 hour) and then Proteinase K (65°C, 2 hours). DNA was purified using the ChIP DNA Clean & Concentrator kit (Zymo Research). Indexed sequencing libraries were generated using the NEBNext Ultra II DNA Library Prep kit (NEB, # E7645), pooled and sequenced on an Illumina HiSeq instrument as paired end 150 bp reads (Genewiz, NJ).

### ChIP-seq analysis

Demultipexed fastq files were mapped to the sacCer3 assembly of the *S. cerevisiae* genome as 2×36mers using Bowtie (v.1.0.1) (Langmead et al., 2009) with the following settings: -v 2 -k 2 -m 1 -- best --strata. Duplicate reads were removed using picard-tools (v.1.99). Peaks were called using MACS2 (v.2.1.0) (Feng et al., 2012) with the following settings: -g 12000000-f BAMPE. RPM (Reads Per Million) normalized read coverage genome browser tracks were generated using custom-written python scripts.

### Cell cycle and protein partitioning modeling

The cell cycle was modeled as reported in (Chandler-Brown et al., 2017). We simulated the entire cell cycle, where cells grew and divided according to measured growth and cell cycle transition rates. This accounts for cell-to-cell variability. To examine the role of protein partitioning in the overall scaling of protein expression, we simulated the synthesis of a constitutively expressed protein (*p*) in each cell (Fig. 5D & S7B). Within the model, protein synthesis and partitioning properties were varied. Protein synthesis was modelled as either scaling in proportion to cell size 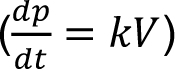 or constant independent of cell size 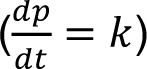 and protein partitioning was modeled as either volume-proportional partitioning at cytokinesis 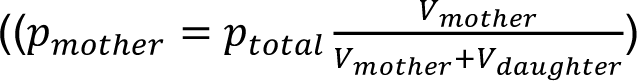 and 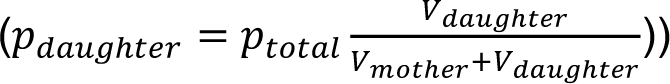 or partitioned in the manner empirically measured for Whi5, where a significant fraction is partitioned by amount 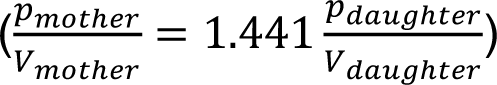. Cells were simulated until a steady-state distribution was achieved and all cells at the last time-point were plotted.

